# Halofilins as Emerging Bactofilin Families of Archaeal Cell Shape Plasticity Orchestrators

**DOI:** 10.1101/2024.01.22.576759

**Authors:** Zachary Curtis, Pedro Escudeiro, John Mallon, Olivia Leland, Theopi Rados, Ashley Dodge, Katherine Andre, Jasmin Kwak, Kun Yun, Berith Isaac, Mar Martinez Pastor, Amy K. Schmid, Mechthild Pohlschroder, Vikram Alva, Alex Bisson

## Abstract

Bactofilins are rigid, non-polar bacterial cytoskeletal filaments that link cellular processes to specific curvatures of the cytoplasmic membrane. Although homologs of bactofilins have been identified in archaea and eukaryotes, functional studies have remained confined to bacterial systems. Here, we characterize representatives of two new families of archaeal bactofilins from the pleomorphic archaeon *Haloferax volcanii*, halofilin A (HalA) and halofilin B (HalB). HalA and HalB polymerize *in vitro*, assembling into straight bundles. HalA polymers are highly dynamic and accumulate at positive membrane curvatures *in vivo*, whereas HalB forms more static foci that localize in areas of local negative curvatures on the outer cell surface. Gene deletions and live-cell imaging show that halofilins are critical in maintaining morphological integrity during shape transition from disk (sessile) to rod (motile). Morphological defects in Δ*halA* result in accumulation of highly positive curvatures in rods but not in disks. Conversely, disk-shaped cells are exclusively affected by *halB* deletion, resulting in flatter cells. Furthermore, while Δ*halA* and Δ*halB* cells imprecisely determine the future division plane, defects arise predominantly during the disk-to-rod shape remodeling. In fact, the deletion of *halA* in the haloarchaeon *Halobacterium salinarum*, whose cells are consistently rod-shaped, impacted morphogenesis but not cell division. Increased levels of halofilins enforced drastic deformations in cells devoid of S-layer, suggesting that HalB polymers are more stable at defective S-layer lattice regions. Our results set halofilins apart from their bacterial correlate, where they provide mechanical scaffolding instead of directing envelope synthesis.

## Introduction

Cell shape maintenance is a universal trait and a focus of interest in every studied organism (1, 2). Cells employ diverse strategies to achieve shape control, such as modulating the composition of the cell wall and the biophysical and biochemical properties of the cytoskeletal polymers that direct their organization (3–5). However, little is known about such mechanisms in Archaea, where most known species lack a rigid, three-dimensional cell wall (6). Instead, the archaeal outer surface of the cytoplasmic membrane is coated by the S-layer, a lattice structure composed of glycoproteins (7, 8). Although the S-layer lattice is assembled through lateral, non-covalent protein-protein interactions, the S-layer glycoprotein (SLG) can be covalently anchored to the cytoplasmic membrane by lipidation in some archaea (9, 10). Ultimately, the S-layer serves as a spatiotemporal reference for the biogenesis of appendage structures, such as pili and archaella (8). Still, its role in cell shape biogenesis remains a topic of debate (11–13).

Haloarchaea are a class of salt-loving microbes notable for their vast morphological diversity, including rods, cocci, triangular, square, and other polygonal-shaped cells (14). Furthermore, the model organism *Hfx. volcanii* has been shown to reversibly shape-shift between two developmental states: motile (rod-shaped cells) and sessile (polygonal, disk-shaped cells) (15–17). Mutations in the archaeosortase ArtA, required for the lipidation of SLG, result in an imbalance in the rod and disk equilibrium, favoring rod-shaped cell formation (9). Moreover, many *Hfx. Volcanii* cell-shape factors have been identified and characterized recently, with cytoskeletal polymers and associated proteins playing a central role. Among them, the tubulin-like paralogs FtsZ1 and FtsZ2 assemble the cytokinetic ring (Z-ring) and couple cell division with shape (18). Another superfamily of tubulin homologs, the CetZ proteins, direct rod-shape formation and motility (16). In contrast, the actin homolog volactin is required for the morphogenesis and viability of disk-shaped cells (19).

Although most microbes regulate cell shape by patterning the cell wall, it is unclear whether archaeal cells control morphogenesis through the turnover of the S-layer lattice or analogous to eukaryotic cell-wall-less cells. Eukaryotic cytoskeleton systems often fulfill multiple functions, such as cargo transport, mechanosensing, and scaffolding (20, 21). Meanwhile, many bacterial polymers could be classified as “cytomotive”, dynamically localizing synthases that build, for example, organized cell wall structures, thereby playing the role of an exoskeleton by mechanically supporting cell shape (22–24). Distinguishing between these two models is particularly relevant for cells undergoing shape transitions, such as *Hfx. volcanii*.

Bactofilins are a class of widely distributed bacterial cytoskeletal polymers with a conserved subdomain featuring a right-handed β-helical fold with highly variable N- and C-terminal ends (25–27). Unlike most cytoskeletal proteins such as actin and tubulin, bactofilins spontaneously self-assemble into non-polar filaments independent of nucleotides and other cofactors (28, 29). *In vitro* studies have shown bactofilin polymers to be mechanically stable and resistant to high salt conditions and chelating agents (28). Their intrinsic membrane affinity and rigidity are crucial for securing the recruitment of associated enzymatic complexes at specific subcellular locales (30, 31). Furthermore, bactofilin filaments have been characterized in many different species, playing essential roles in a multitude of cellular processes, including cell shape and stalk maintenance (31–35), chromosome segregation (36), and cell polarity (37).

Until recently, bactofilins were considered exclusive to bacteria. However, work by Deng and colleagues has expanded the bactofilin universe to eukaryotes and archaea (29). Here, we use computational structural prediction, reverse genetics, and single-cell microscopy to characterize the evolutionary organization and *in vivo* function of two new archaeal bactofilins, halofilin A (HalA) and halofilin B (HalB).

## Results

### Halofilins represent diverse families of the bactofilin superfamily

A sequence search for bactofilin homologs in the archaeal model *Hfx. volcanii*, using the Pfam bactofilin domain (PF04519) as query, yielded two previously reported candidates, now designated halofilin A (HVO_1610 / UniProtKB D4GZ39) and halofilin B (HVO_1237 / Uni-ProtKB D4GX31). Subsequent analysis of the structures predicted by AlphaFold2 (38) revealed that HalA consists of two bactofilin-like domains arranged head-to-head and connected by an α-helical hairpin (between residues Arg122 and Asp161). The N- and C-termini of HalA exhibit disordered tails (Fig. 1A), a feature also common in bactofilins. Each bactofilin subdomain in HalA has six right-handed windings of parallel β-strands that form a single-stranded, three-faced, right-handed β-helix, equivalent to canonical bactofilins. Notably, the N- and C-terminal bactofilin subdomains of HalA superimpose at a root-mean-square deviation (RMSD) of ∼1 Å and superimpose with *Thermus thermophilus* bactofilin (TtBac) at RMSDs of ∼2 Å and ∼1.6 Å, respectively. To identify potential polymerization interfaces of HalA, we used AlphaFold2-Multimer (38, 39) to generate homodimer predictions. In most of these models, the HalA monomers are arranged C-terminus-to-C-terminus (Fig. S1D; pTM = 0.61, ipTM = 0.34), with no cases of N-terminus-to-N-terminus observed (Fig. S1A-D). Interestingly, in one of the top-ranking models, the dimerization interface is formed by the α-helical hairpins (Fig. S1E; pTM = 0.49, ipTM = 0.1299). The first α-helix of the hairpin motif is enriched in hydrophobic amino acids (Fig. S1F) and could also serve as a membrane-targeting sequence (MTS). We also attempted to predict structures for homotrimers and homotetramers of HalA using AlphaFold2-Multimer; however, in the predicted oligomeric structures, the monomers did not arrange into filaments. As a control, we used AlphaFold2-Multimer to predict homo-oligomers of TtBac, obtaining filaments with subunits arranged in a head-to-head configuration, which is in agreement with the solved TtBac filament structure.

**Figure 1.**
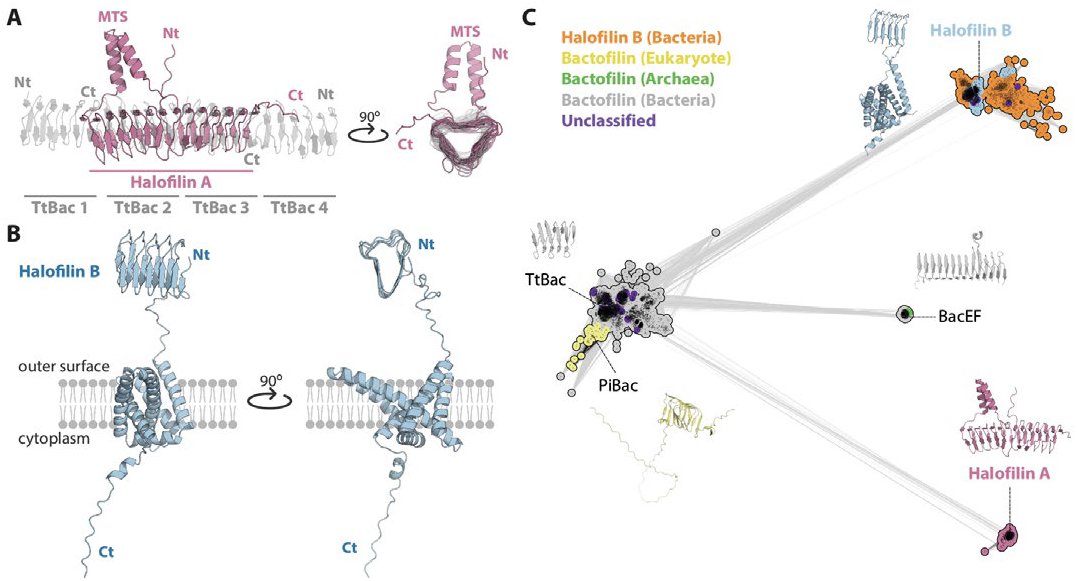
Halofilins comprise two large subgroups of structurally and evolutionarily diverse bactofilin homologs. **(A)** Predicted structure of HalA from Hfx. volcanii. A HalA monomer is overlayed to a TtBac filament with three subunits (PDB entry 6RIA). MTS = membrane-targeting sequence. **(B)** Membrane positioning of the predicted structure of HalB from Hfx. volcanii. The structure is lacking the signal peptide (see Methods). **(C)** Cluster map of canonical bactofilins together with HalA and HalB. A set of 8,542 protein sequences is shown. Each dot represents one protein sequence, and each line represents pairwise similarity between two sequences, as calculated by BLASTp. Representative sequences are annotated for each cluster, and a cartoon representation of their predicted structure except for TtBac, solved experimentally (PDB 6RIA-A) - is shown next to it. Clustering was achieved at an E-value cutoff of 1e-10, and edges in the map are shown for E-values ≤ 1e-8. Seed sequence and structure model accessions are provided in Tables S1 and S2.

Unlike HalA, HalB is predicted to be an α-helical integral membrane protein with a Sec/SPI signal peptide and five transmembrane (TM) helices. A sixth, shorter helix is positioned parallel to the inner leaflet of the membrane (Fig. 1B). The very high (>90) pLDDT values suggest that this TM fold was modeled to high accuracy (Fig. S2A-B). In addition, HalB has an extracellular bactofilin domain with five windings at its N-terminus (residues 41-125, henceforth denoted as bHalB), contrasting the six found in HalA and TtBac, and possesses a disordered cytoplasmic tail at its C-terminus. The Predicted Aligned Error (PAE) values are low throughout the entire putative structure of HalB (Fig. S2C), indicating that AlphaFold2 predicts well-defined relative positions and orientations for all residues. The bactofilin domain of HalB superimposes with the bactofilin subdomains of HalA at RMSDs of ∼2 Å and with TtBac at ∼1.5 Å. Notably, AlphaFold2-Multimer models for homo-oligomers of bHalB also did not yield conclusive oligomers, likely being spurious predictions.

To systematically assess the relationship between HalA, HalB, and canonical bactofilins at the sequence level, we performed pairwise BLASTp comparisons of (i) HalA and TtBac, (ii) bHalB and TtBac, and (iii) HalA and bHalB. The C-terminal bactofilin subdomain of HalA (residues 180-259) shows apparent homology to TtBac (identity = 29.79%, query coverage = 72%, E-value = 5e-6), whereas the N-terminal subdomain aligns only partially and with very low significance (E-value = 0.47) to TtBac. To further investigate the evolutionary relationship between the N-terminal bactofilin subdomain of HalA and canonical bactofilins, we used the sensitive remote homology detection method HHpred, which is based on profile Hidden Markov Models (HMMs). Through this analysis, we found a statistically significant match between the N-terminal subdomain of HalA (residues 12-114) and the Pfam bactofilin domain PF04519 (Pr = 89.39%, E-value = 0.043). These results indicate that the N-terminal subdomain shows greater divergence from canonical bactofilins compared to the C-terminal subdomain. BLASTp found no statistically significant similarity between bHalB and either TtBac or HalA. However, using HHpred, we obtained statistically significant matches between bHalB and TtBac (Pr = 98.53%, E-value = 1.8e-15) and between bHalB and the C-terminal bactofilin subdomain of HalA (Pr = 93.76%, E-value = 1.8e-7), suggesting that bHalB is also homologous to bactofilins.

To further evaluate the structural and sequence similarity between bactofilins and halofilins, we aligned the predicted AlphaFold2 structures for the N- and C-terminal bactofilin domains of HalA, bHalB, and those of 14 canonical bactofilins (Fig. S3A). The domains superimpose with an average TM-score of 0.86, an average RMSD of 1.21 Å, and an average aligned length of 92 residues, suggesting very close structural similarity. We also used the respective sequences from this structural alignment to create a structure-guided and manually curated multiple sequence alignment (MSA) (Fig. S3B). The sequence conservation is pronounced particularly the preservation of Glycines in the turns, and of hydrophobic residues in the β-strands, consistent with earlier findings (40, 41).

To substantiate the observed sequence relationship between HalA, HalB, and bactofilins on a global scale, we assembled their homologs from UniProtKB and clustered them using CLANS (42) based on the strength of their all-against-all pairwise sequence similarities (see Methods for details). In the resulting cluster map, HalA, HalB, and bactofilin sequences are organized into three distinct clusters (Fig. 1C). At the E-value cut-off (1e-8) chosen to visualize the connections, the HalA and HalB clusters are connected to the bactofilin cluster but not to each other, indicating a closer relationship between HalA and bactofilins. Eukaryotic bactofilins, found in stramenopiles and ascomycetes and exemplified in the cluster map by the stramenopile *Phytophthora infestans* bactofilin PiBac, cluster tightly together with canonical bacterial bactofilins. This supports the hypothesis that these eukaryotes may have acquired bactofilins from bacteria by horizontal transfer, as previously proposed (29). Besides the HalA, HalB, and bactofilin assemblages, our map contains a fourth, smaller group radiating from the bactofilin cluster (BacEF). This cluster consists of bacterial and archaeal bactofilin-like proteins with 13 windings, represented by homologs of BacEF (UniProtKB P39132 and P39133) from *Bacillus subtilis* (30, 43) and the uncharacterized protein from *Methanobacterium bryantii* (UniProtKB A0A2A2H537).

The HalA cluster consists of 681 proteins exclusively from Archaea (Fig. S4), with most homologs originating from the phylum Euryarchaeota, particularly from Halobacteria (n = 382) and Methanomicrobia (n = 192). Additionally, we found some homologs in archaeal candidates and unclassified phyla as well as in the Asgard group. Analysis of AlphaFold2-predicted structures for numerous HalA sequences revealed that although they share a head-to-head stacked fold, an equal number of windings in each bactofilin subdomain, and the α-helical hairpin, they exhibit great diversity in their N- and C-terminal α-helices and disordered tails (Fig. S6).

In contrast, the HalB cluster, consisting of 2,304 proteins, shows a broader distribution across both Archaea and Bacteria, with a higher prevalence (n = 1,813) in the latter (Fig. S5). Similar to HalA, archaeal HalB is predominantly found in Euryarchaeota (n = 454), especially in Halobacteria (n = 280) and Methanomicrobia (n = 152). Within Bacteria, HalB is most abundant in Chloroflexota (n = 387), Pseudomonadota (n = 287), and Bacillota (n = 170). Additionally, HalB is found in numerous unclassified clades, candidate phyla, and divisions of both Bacteria and Archaea. HalB variants also exhibit broader structural diversity (Fig. S6). For example, the bHalB domains comprise up to ten windings among all the analyzed structures. The C-terminal tail also shows significant variability; in some cases, it is associated with an additional domain and may extend further into the cytoplasm (e.g., A0A3B9P870), while in others it is absent (e.g., A0A662QX32). Nonetheless, all HalB homologs have a TM domain with the same number of □-helices preceded by an unstructured segment.

The widespread taxonomic distribution of bactofilins across the three domains of life suggests that a primordial form of these proteins may have existed at the time of the last universal common ancestor (LUCA). It is likely that HalA originated from the duplication, fusion, and subsequent diversification of an ancestral bactofilin-like gene. Similarly, HalB might have arisen by the fusion of an ancestral bactofilin-like gene with one encoding a TM domain, followed by diversification (Fig. S6). Notably, HalA and HalB are predominantly found in the Halobacteria and Methanomicrobia classes (Fig. S4 and S5), indicating the possibility of an underlying co-evolutionary relationship between these proteins.

### Halofilin mutants are dominant-negative in distinct cell types

To investigate the function of halofilins, we created strains deleted of *halA, halB*, and the double mutant Δ*halAB*. We started by characterizing standard microbiological phenotypes. Although mutant strains showed no significant differences in bulk growth and viability (Fig. S7A-B), all three strains showed a drastic decrease in motility in comparison to the wild type (51±9%, 79±10%, and 38%±11%, respectively) (Fig. S7C). Phenotypes of *Hfx. volcanii* mutants often include confounding factors due to physiological differences under different growth stages. Many haloarchaeal species exhibit developmental shape transitions between disks (pili-based, sessile cells) and rods (archaella-based swimming cells). Shape transitions have been shown to occur across bulk growth phases, with the majority of rods or disks found in the early-exponential and mid-exponential phases, respectively (15, 17). Consequently, the motility defects of halofilin mutants could be related to rod shape.

To determine how halofilin deletions impact different cell types, we initially imaged cultures with mixed populations of rods and disks using live-cell super-resolution microscopy (Fig. 2A, Movie S1) combined with phase contrast microscopy (Fig. S8A). To compare cell shapes from different strains using a consistent metric across cell types, we measured cell solidity (S), which denotes how much a cell outline stray away from a perfect solid object on a normalized scale from 0 (most deformed) to 1 (most regular) (44). Using wild-type cells as reference (S=0.90±0.02 for rods and S=0.98±0.02 for disks), the deletion of *halA* resulted in elongated, irregular rods (S=0.82±0.04), while disks were only mildly affected (S=0.89±0.03) (Fig. 2B and S8B). Nonetheless, Δ*halB* rods were indistinguishable from wild type (S=0.89±0.03), but Δ*halB* disks appeared larger and more angular (S=0.84±0.04). Moreover, rods of the Δ*halAB* double mutant combine the cell shape defects of both Δ*halA* rods (S=0.81±0.03) and Δ*halB* disks (S=0.83±0.05). Therefore, the rod-related morphology phenotypes in Δ*halA* and Δ*halAB* correlate with the observed motility deficiencies (Fig. S7C). These results suggest that HalA and HalB may coordinate non-redundant functions or be relevant in different lifestyles of *Hfx. volcanii* cells.

**Figure 2.**
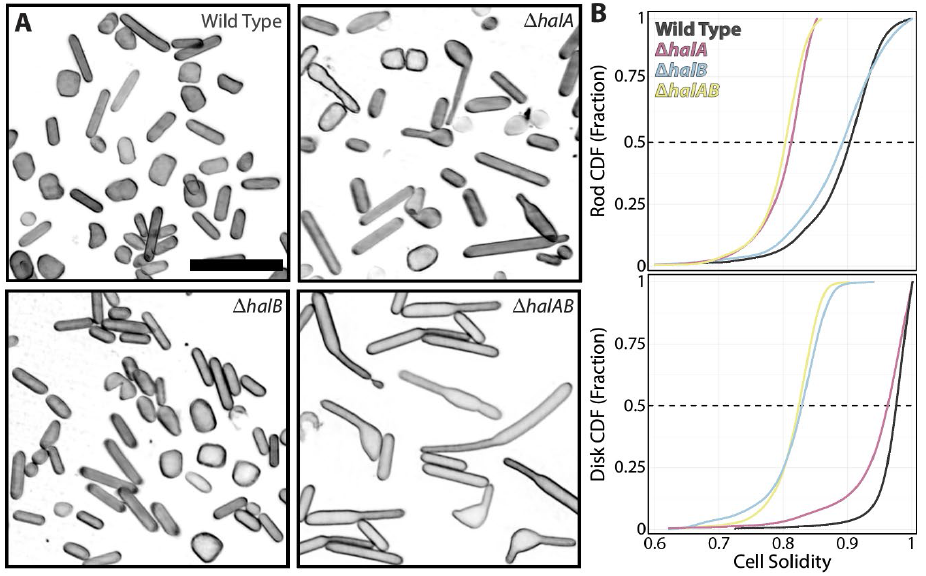
HalA and HalB play specific roles in rods and disks, respectively. **(A)** 3D-SoRa super-resolution projections of wild-type cells compared to deletions of halA, halB, and halAB. Cell membranes were labeled with Nile Blue. **(B)** Cumulative distributions of rods (top) and disks (bottom) cell solidities from phase contrast images. Dashed lines indicate the median values of the distribution. Kolmogorov-Smirnov tests were applied to pairs between every dataset, resulting in statistically significant differences (p<0.0001) as detailed in Figure S8.

### Halofilins are bona fide cytoskeletal proteins

To determine whether HalA and HalB are genuine cytoskeletal proteins that control cell shape by polymerizing in their native cellular environment, we created merodiploid strains to ectopically express C-terminal msfGFP fusions. Imaging cells using epifluorescence microscopy, HalA-msfGFP was found to assemble into relatively long filaments across the major axis of cells (Fig. 3A, top left panel). 3D projections from super-resolution images revealed curved HalA filaments on the inner surface of the cytoplasmic membrane (Fig. 3A, top right panel, Movie S2). In contrast, HalB-msfGFP is distributed as bright foci distributed around the cell’s contour (Fig. 3A, bottom panels, Movie S2). Interestingly, other than a few subtly elongated patches, we did not observe HalB-msfGFP forming pronounced polymer structures akin to HalA-msfGFP. Importantly, the ectopic expression of both fusions rescued cell shape defects when complementing their respective knockout strains (Fig. S8), suggesting these structures should be physiological.

**Figure 3.**
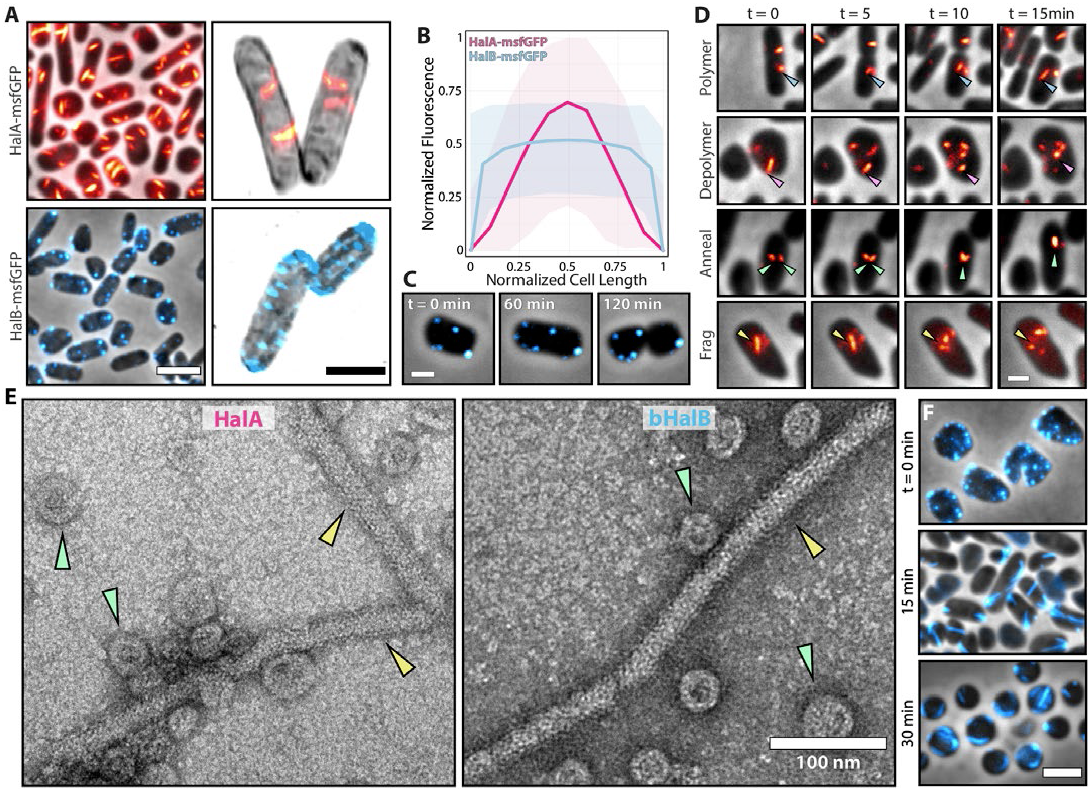
HalA assembles into dynamic filaments and HalB as distinct foci in live cells. **(A)** Widefield fluorescence and phase contrast (left column) and 3D-SoRa super-resolution (right column) micrographs of cells expressing HalA-msfGFP (orange) and HalB-msfGFP (blue). Membranes of cells from 3D-SoRa images were stained with Nile Blue (grey). **(B)** Quantitation of the relative localization of HalA-msfGFP (pink) and HalB-msfGFP (blue). Phase-contrast images of cells were segmented, and the coordinates of the major axis of the cells were correlated to the mean grey value of the msfGFP channel at 0.5-pixel level. The length of the major axis and the msfGFP fluorescence were normalized for each cell out of three biological replicates. **(C)** Phase-contrast images and epifluorescence microscopy time-lapse of HalB-msfGFP assembles as slow-diffusing foci across the cell membrane. **(D)** Phase-contrast and epifluorescence microscopy time-lapses of HalA-msfGFP cells show a plethora of dynamic behaviors from HalA polymers. **(E)** HalA and bHalB polymerize in vitro. Transmission Electron Microscopy (TEM) of purified HalA and bHalB polymers showing filament bundle (yellow arrowheads) and toroid (green arrowheads) structures. **(F)** Phase-contrast images and epifluorescence microscopy time-lapse of HalB-msfGFP extensive polymerization following spheroplasting by 50 mM EDTA. Unless specified, scale bars represent 2 µm.

To characterize the localization of each halofilin fusion in the cell, we plotted the fluorescence intensity profile from hundreds of cells under the microscope against their position across the major cell axis (Fig. 3B). While HalB-msfGFP foci do not appear to populate specific cellular locales, HalA-msfGFP filaments are enriched around the midcell vicinity.

Next, we collected time lapses with epifluorescence across multiple generations to assess whether HalA and HalB structures are dynamically regulated throughout the cell cycle. Like BacEF bactofilins that orchestrate flagella placement (30, 43), HalB foci are relatively static across the scale of minutes (Fig. 3C, Movie S3). Conversely, HalA polymers are unusually more dynamic than the studied bactofilins, exhibiting a plethora of polymerization behaviors (Fig. 3D, Movie S4). The contrast between HalA and HalB localization and dynamic profiles suggest they may only interact transiently, if at all. HalA and HalB localization and dynamic profiles suggest they may only interact transiently, if at all. The possible decoupling between HalA and HalB aligns with their non-overlapping functions, as suggested by the specific phenotypes from deletion strains.

To test if HalA polymers assemble independently of other known polymer systems, we expressed HalA-msfGFP and HalB-msfGFP in cells carrying deletions of every known *Hfx. volcanii* cytoskeleton genes (including Δ*halA* and Δ*halB*): Δ*ftsZ1*Δ*ftsZ2* (18), Δ*cetZ1* to Δ*cetZ6* (16), and Δ*volA* (19). In all cases, HalA-msfGFP and HalB-msfGFP assembled into filaments and foci, respectively (Fig. S9A). Curiously, we observed changes in HalA-msfGFP length and number of HalB foci in all strains (Fig. S9B). However, the non-specific nature of these perturbations to HalA and HalB assembly profiles suggests that variations are likely caused by shape defects rather than direct interactions between these polymer systems.

To determine if halofilins self-associate independently of any cellular context, we heterologously expressed and purified HalA and bHalB (residues Gly36-Asp127) from *Escherichia coli*. Interestingly, both HalA and bHalB protein fractions showed bands higher than expected for their sizes (approximately 30 and 8 kDa, respectively) in gels (Fig. S10A). This phenomenon has been observed before with proteins from halophilic organisms, likely due to glycosylation or high negative surface charge that could drastically influence their migration in SDS (45, 46). We then imaged purified fractions by negative staining transmission electron microscopy (TEM). From both HalA and bHalB samples, we consistently observed straight filament bundles (yellow arrowheads) and toroid structures (green arrowheads) (Fig. 3E). The HalA straight bundles were significantly shorter than those of HalA-msfGFP observed in cells (186±53 nm and 570±62 nm, respectively) (Fig. S10B). In contrast, bHalB bundles were larger than those of HalA (489±294 nm), large enough to be resolved *in vivo* as well (Fig. S10B). Curiously, the HalA and bHalB bundles are similar to the micrographs by Zuckerman *et. al* (25) showing bactofilin BacM polymers *in vitro*. As a control, we imaged the fractions from non-induced cultures, and did not observe large bundles but only a few (larger) toroid structures in the induced samples (Fig. S10C).

Finally, we sought to understand why HalB would not assemble into larger polymeric structures in vivo. Based on our structural predictions (Fig. 1B), we noted that bHalB may be placed approximately 75Å above the membrane outer surface and close to the S-layer lattice plane (47). Therefore, we speculated that the S-layer lattice could be directly or indirectly inhibiting the polymerization of HalB in cells. To test this hypothesis, we treated cells expressing HalB-msfGFP with 50 mM EDTA to disrupt the S-layer lattice by chelating Ca^2+^ ions (48). During the spheroplasting process, even before cells lost their shape and became spherical, we observed long HalB-msfGFP filaments emerging from the initial foci, possibly from places where the S-layer lattice was initially disassembled, persisting throughout the spheroplasting course (Fig. 3F).

### bHalB bactofilin subdomain is located at the extracellular surface

HalB represents the first multipass transmembrane protein with its bactofilin domain predicted to be cell-surface exposed, distinguishing it from other bactofilins that are cytoplasmic. Based on our structural predictions and the HalB-msfGFP spheroplasting experiment, we reasoned that the bHalB position relative to the S-layer may play a role regulating its polymerization. First, we confirmed that HalB is in fact a membrane protein by western blots using anti-GFP antibodies of cells expressing HalB-msfGFP (membrane) and free msfGFP (cytoplasm). As expected, HalB-msfGFP bands were observed only in pellets, while free msfGFP was found in the supernatant (Fig. S11A).

Next, to validate our model of HalB membrane topology, we combined HalB fusions with HaloTag and the (non)permeability properties of Alexa488 and JF549 to cell membranes. While HaloTag technology has been widely employed *in vivo* across many bacteria and in the haloarchaeon *Halobacterium salinarum* (36), some microbes, such as fission and budding yeast, require transporter deletions to retain the dyes in the cell (49, 50). To validate the HaloTag in *Hfx. volcanii*, we incubated cells expressing cytoplasmic HaloTag with a mix of the probes JF549 (permeable) and Alexa488 (non-permeating) HaloTag ligands. Consequently, we observed a cytoplasmic HaloTag-JF549 signal but not Alexa488 (Fig. S11B, top row). As a control, we also incubated wild-type cells (without HaloTag expression) and observed no significant labeling with either JF549 or Alexa488 HaloTag ligands (Fig. S11B, bottom row). Based on these observations, we concluded that HaloTag is a suitable reporter for specific protein labeling in live *Hfx. volcanii* cells.

To test the HalB topology, we created two new HalB-HaloTag constructs: one mimicking the predicted cytoplasmic C-terminal msfGFP fusion shown in Figure 3, and the second a sandwich fusion inserting HaloTag adjacent to the Ser34 residue between the predicted Sec secretion signal (residues 1-30) and the bHalB subdomain (Fig. 4A, top row). As expected, both HalB-HaloTag constructs were labeled with the cell-permeable JF549, while Alexa488 specifically decorated the extracellular HaloTag fusion (Fig. 4A, bottom row). From the localization, dynamics, and topology profile, we conclude that HalB is a transmembrane protein whose bactofilin domain is exposed on the outer cell surface, linked by a flexible linker long enough to be oriented toward the S-layer plane.

**Figure 4.**
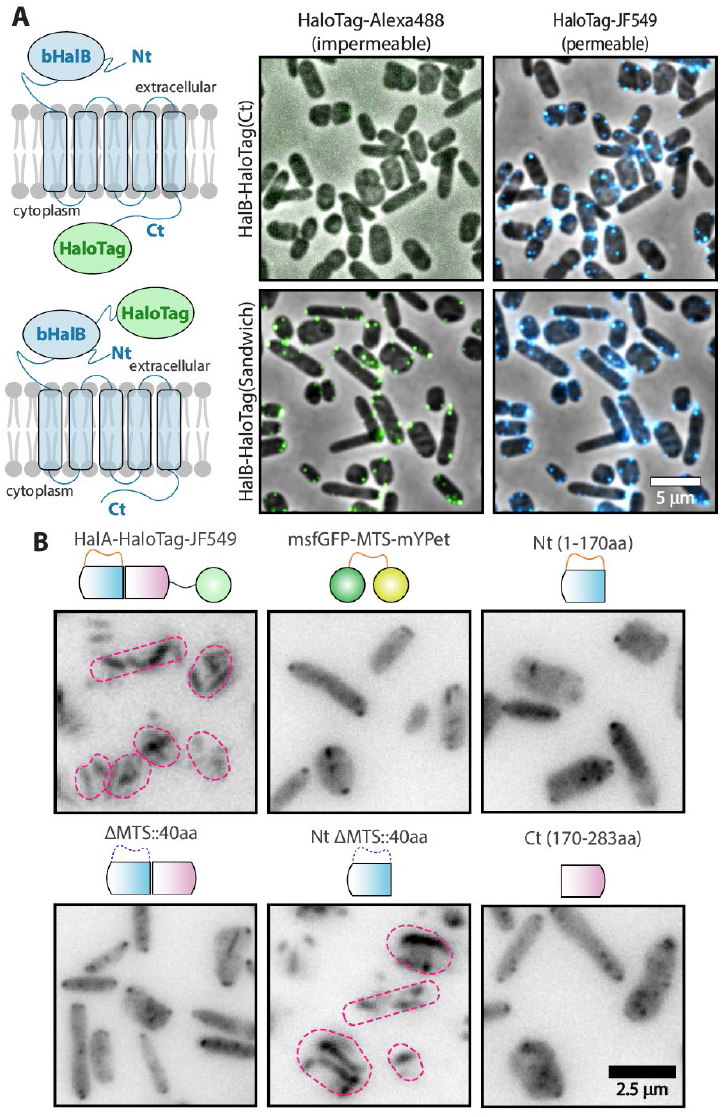
Experimental validation of HalA polymerization and HalB structural topology. **(A)** Cartoons representing the HalB-HaloTag fusions created by tagging HalB at its C-terminal (predicted cytoplasmic region, top panel) or next to its bactofilin subdomain (predicted extracellular region, bottom panel). Cells expressing the HalB-HaloTag fusions were incubated simultaneously with 100 nM of Halo-Tag-ligand Alexa488 (left panels, green) and JF549 (right panels, blue) and imaged by phase contrast and epifluorescence microscopy. Scale bar represent 5µm. **(B)** Cells co-expressing HalA-HaloTag together with different truncated versions of HalA. Depolymerization of HalA was extensive in most constructs but under the N-terminal subdomain alone (bottom-center panel). Cell outlines were drawn to support the identification of HalA polymers.

### The MTS and the C-terminal subdomain are required for HalA polymerization

To further understand the structural principles underlying HalA polymerization, we created a series of truncated HalA-msfGFP constructs and tested whether they could perturb HalA filaments tagged with HaloTag. Since HalA-HaloTag assembled into polymers like HalA-msfGFP (Fig. 4B, top-left panel), we proceeded to test if the co-expression of the selected constructs would disturb the polymerization of HalA. First, we tested whether the MTS region is required for HalA polymerization by replacing the MTS (R122-Asp161) with a 40 amino acid-long flexible linker (ΔMTS::40aa). Expression of HalA lacking MTS extensively depolymerized the HalA filaments, suggesting that the HalA(ΔMTS::40aa) can still be incorporated into polymers but with decreased affinity for the membrane (Fig. 4B, top-center panel). A non-mutually exclusive hypothesis is that the MTS sequences may be involved in the polymerization contact of HalA filaments (Fig. S1E). To probe the MTS interaction with the HalA polymers, we expressed HalA(MTS) alone, positioned between msfGFP and mYPet fluorescent proteins. This “sandwich” construct was created in such way to mimic the MTS intercalation between HalA’s Nt and Ct and to avoid unfolded regions that could be targeted for degradation. Like the HalA(ΔMTS::40aa) construct, the MTS was sufficient to depolymerize HalA, suggesting that the α-helical hairpin may be involved in more than just HalA membrane anchoring (Fig. 4B, top-right panel).

Next, we investigated the role of the N-terminal and C-terminal ends of HalA and whether they are, as in canonical bactofilins, part of the polymerization surface. For this purpose, we expressed the N-terminal subdomain (1-170aa) and observed extensive depolymerization of HalA (Fig. 4B, bottom-left panel). However, this region encompasses the MTS, which alone can depolymerize HalA. Hence, we created an N-terminal subdomain in which the MTS region was replaced with a 40aa flexible linker. This time, unlike bacterial bactofilins, the N-terminal subdomain lacking the MTS was insufficient to depolymerize HalA filaments (Fig. 4B, bottom-center panel). Finally, the expression of the C-terminal sub-domain alone induced depolymerization of HalA (Fig. 4B, bottom-right panel).

To confirm that protein levels, stability and predicted localization of the truncate fusions are as expected for functional probes, we probed the cellular localization of the truncate constructs expressed in Δ*halA* cells (Fig. S11C). As expected, the full-length HalA-msfGFP assembled filaments as in the merodiploid strain. Moreover, both MTS constructs (msfGFP-MTS-mYPet and Nt) localized around cell outlines independently of HalA filaments, supporting the affinity of the MTS region to membrane. Likewise, the last three constructs lacking the MTS region (ΔMTS, Nt-ΔMTS and Ct) showed dispersed signal in the cytoplasm. Importantly, none of these constructs showed evidence of aggregation. Quantification of the fluorescence signal from cells indicated that all fusions are being expressed at similar levels relatively to each other, and higher levels compared to HalA-msfGFP (Fig. S11D).

Finally, we verified if the (cytoplasmic) Nt-ΔMTS truncate construct is intact, or undergoing proteolysis and releasing msfGFP in the cytoplasm. To investigate this scenario, we performed Western blots using anti-GFP antibodies from HalA and Nt-ΔMTS extracts. For sample load normalization, we co-expressed free msfGFP under the control of a constitutive promoter (P250) recently characterized by our group (57). In agreement with our microscopy measurements, HalA and Nt-ΔMTS showed comparable concentrations (HalA-msfGFP 10% higher than HalA-Nt) and no obvious evidence of proteolysis (Fig. S11E). Altogether, these results suggest that the MTS and C-terminal regions, but not the N-terminus, play an important role in the polymerization, membrane interaction, and/or stability of HalA filaments.

### Halofilin mutants indirectly cause decondensation of cytokinetic rings

Based on the accumulation of HalA-msfGFP around the midcell position (Fig. 3B) and morphological defects observed at the cell poles of halofilin mutant cells (Fig. 2A), we sought to investigate whether HalA and HalB orchestrate cell division. Most archaea assemble their cell division machinery by assembling the Z-ring with two tubulin-like cytoskeletal filaments, FtsZ1 and FtsZ2. Recently, Liao *et al*. proposed that each FtsZ paralog might play non-overlapping roles, with FtsZ1 involved in cell elongation/shape and FtsZ2 in cytokinesis (18). To probe the Z-ring localization in the halofilins mutants, we expressed FtsZ1-GFP and FtsZ2-GFP in the background of each mutant strain and imaged rod and disk cell populations (Fig. 5A-B and S12A-B). We found that Z-rings decorated with either FtsZ1-GFP or FtsZ2-GFP showed greater positional variance across the long axis of the halofilin mutant rod cells, with a more pronounced decondensation of Z-rings in the Δ*halA* (σ^2^=0.45 and 0.47, respectively) and Δ*halAB* (σ^2^=0.42 and 0.45, respectively) mutants compared to wild type (σ^2^=0.36 and 0.33, respectively) and Δ*halB* (σ^2^=0.38 and 0.35, respectively) (Fig. S12A). In contrast, we did not observe as drastic a decondensation of the Z-rings labeled with FtsZ1-GFP across mutant strains when cells were disk-shaped (σ2=0.29 for wild type, 0.33 for ΔhalA, 0.35 for Δ*halB* and 0.39 for Δ*halAB*). However, we observed consistent decondensation for Z-rings labeled with FtsZ2-GFP (σ2=0.18 for wild type, 0.22 for Δ*halA*, 0.38 for Δ*halB* and 0.39 for Δ*halAB*) (Fig. S12B). The differences between FtsZ1 and FtsZ2 fusions could be explained by a possible difference in functionality of each probe in disks but not in rods, although both fusions seem to localize similarly in the wild type background.

**Figure 5.**
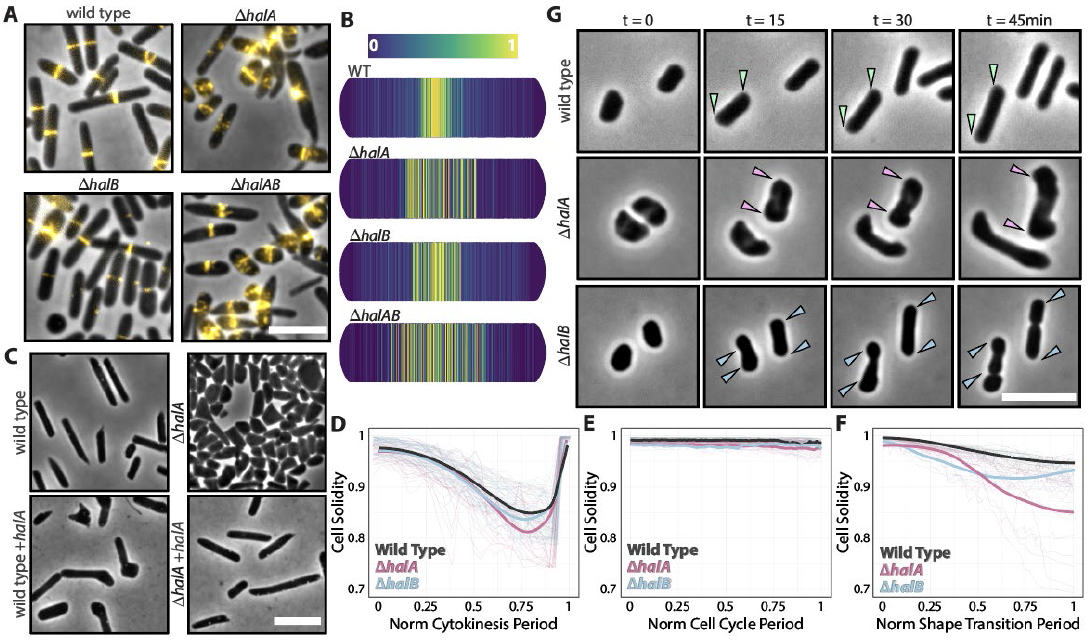
ΔhalA and ΔhalB mutant cells show Z-ring decondensation and morphological defects during shapeshift. **(A)** Phase-contrast and epifluorescence microscopy of cells expressing FtsZ2-msfGFP (labeling the cytokinetic Z-ring). **(B)** Normalized heatmaps of the relative localization of the FtsZ2-msfGFP intensity profiles across the major axis of rod cells in wild-type and mutant populations. **(C)** Phase-contrast of Hbt. salinarum wild-type and ΔhalA cells. Deletion (top-right) and overexpression (bottom-left) of halA caused shape-related phenotypes, but not cell division defects. Ectopic expression of halA significantly rescued the morphology of ΔhalA cells (bottom-right), supported by quantitation in Figure S12C. **(D)** Single-cell tracking of cell solidity plotted against normalized cytoki-nesis periods depicting the deformation profile of cells during cell division. A Lorentzian function was used to fit the deformations (constriction) and relief (separation) events from the wild-type (29 cells, RMS error 0.01), ΔhalA (31 cells, RMS error 0.05), and ΔhalB (28 cells, RMS error 0.03) strains. **(E)** Single-cell tracking of cell solidities plotted against a normalized cell cycle period depicting the deformation profile of non-transitioning cell elongation during a single doublingtime growth period. A simple average trace across all data points was used to fit the deformations events from the wild-type (32 cells, S=0.99±0.05), ΔhalA (36 cells, S=0.96±0.15), and ΔhalB (29 cells, S=0.98±0.11) strains. **(F)** Single-cell tracking of cell solidities plotted against normalized shapeshift periods depicting the deformation of cells during disk-rod transitions. A Lorentzian function was used to fit the deformations (disk-rod transition) events from the wild-type (40 cells, RMS error 0.02), ΔhalA (41 cells, RMS error 0.09), and ΔhalB (38 cells, RMS error 0.04) strains. **(G)** Representative phase-contrast time-lapses of cells shapeshifting from disks to rods throughout approximately a quarter of the cell cycle period. Arrowheads indicate the cellular regions under shape-shift remodeling. Scale bars represent 5µm.

Although Z-ring condensation is required for bacterial survival (51, 52) and may explain why rod-shaped Δ*halA* and Δ*halAB* mutants have a larger cell area, cell division defects alone cannot explain the Δ*halB* and Δ*halAB* phenotype in disks. It is often challenging to decouple morphological defects caused by cell division from cell-shape defects that cause pleiotropic cell division defects (53, 54). To decouple cell shape and division processes, we studied the role of halofilins in *Hbt. salinarum*, a haloarchaeon that is unable to shape-shift, consistently maintaining its rod shape (14). It has been previously shown that, unlike *Hfx. volcanii, Hbt. salinarum* mutants that specifically impact cell division undergo filamentation but exhibit limited cell shape defects (56). Therefore, we reasoned that by deleting halofilin in this species, we should be able to identify whether halofilins regulate cell division (cell length) or morphogenesis (cell solidity). Using our dataset of halofilin homologs, we identified only one putative HalA (VNG_0507C, UniProtKB Q9HRW9), but no evidence of HalB or any other bactofilin domain-containing proteins. While deletion of *halA* in *Hbt. salinarum* only subtly decreased cell area compared to wild type (3.4±1.4 3.9±1.3 µm^2^, respectively), mutants were no longer rods, assuming polygonal and round shapes resembling *Hfx. volcanii* disks (Fig. 5C and S12C). Ectopic expression of *halA* rescued both size and rod morphology to wild-type levels, and the expression of a second *halA* copy resulted in larger, deformed rods (Fig. 5C and S12C). Extensive cell shape deviations but limited filamentation supports a role for halofilins in morphogenesis as observed in *Hfx. volcanii*.

Next, to directly observe the relationship between cell shape defects and division in *Hfx. volcanii*, we tested whether solidity deviations observed in Δ*halA* (rods) and Δ*halB* (disks) emerged during cytokinesis, which could be driven by Z-ring decondensation. Therefore, we tracked the solidity of healthy single cells from the onset of constriction to complete daughter cell separation. In wild-type cells, solidity should drastically drop during cytokinesis, recovering in a fast, one-step phase upon daughter cell split. If cell constriction creates misshapen cells, we expect to see mutant cells failing to recover their solidity to initial levels. Conversely, we observed that the Δ*halA* and Δ*halB* mutants recovered their solidity to a similar level as the wild type (Figs. 5D and S12D, Movie S5). Interestingly, the Δ*halA* rods showed a steeper increase in solidity than the wild type and Δ*halB*, although the cells recovered solidity levels above 0.9.

Finally, as a control, we tested if the delocalization of FtsZ1-msfGFP in halofilin mutant cells could contribute to cell shape defects by affecting the proper shape maintenance during single-cell elongation (9, 18). Confirming the results from *Hbt. salinarum*, the solidity tracks of Δ*halA* and Δ*halB* cells showed no significant cell shape defects beyond those seen in the wild type (Fig. 5E and S12E, Movie S5). Combining both analyses, we conclude that halofilins are not directly involved in shape maintenance during cell division and elongation.

### Halofilins prevent morphological defects during shape transitions

Most of the haloarchaeal cell shape factors identified in recent years were found to stabilize cell-type morphologies by shifting the apparent ratio of cell shapes across populations to be dominated by rods or disks (9, 16, 19). However, it is not clear whether these regulators play a specific role during shape transitions. Since halofilin mutants still form rods and disks, albeit with partial, cell-type specific defects, we propose that they play a role during the shape transition process.

Among technical challenges, the mechanistic details during shape transitions have so far been limited by the ability to observe the cells shifting under the microscope in real time. Therefore, we developed a protocol to increase the frequency of multiple simultaneous shape transitions, establishing an easy and affordable assay to quantify time and morphological integrity during transitions. Because cells from isolated colonies on plates are mostly disk-shaped and rods emerge very early in exponential growth in liquid, we speculated that these disks from plates should shape-shift in liquid culture as soon as they leave the lag phase. Hence, we streaked fresh plates until colonies were barely visible, resuspended them in liquid media, and moved samples directly to the microscope. As expected, cells started to transition approximately one generation after resumption of growth.

Next, we tracked the solidity of cells shifting from disks to rods. Following the same logic as in the cell division assay, bactofilin mutants should deform during disk-to-rod shape transitions beyond the wild-type control. For tracking, we selected only disk cells with a standard cell shape (solidity>0.95). While wild-type disks transitioned smoothly to rods with mean solidities of approximately 0.95 (Fig. 5F-G, Movie S6), Δ*halA* cells abruptly deformed only during the second half of the transition period, resulting in rods with mean solidities of approximately 0.85. Meanwhile, Δ*halB* cells followed the same trend as Δ*halA*, but with an early solidity decay in the first half of the process, followed by a subtle recovery to mean solidities of approximately 0.93. In summary, halofilin mutants result in more drastic cell shape defects compared to our previous bulk measurements. This suggests that cells rely on halofilins, directly or indirectly, for mechanical support during disk-to-rod shape remodeling. While HalB could prevent deformations in the initial stage of the process when the cells are still disks, HalA would play a role in a later phase, when the cells have almost fully developed into complete rods.

### HalA and HalB bind to specific membrane curvatures

Previous work on *Helicobacter pylori* has shown that the bactofilin CcmA counterbalances MreB-directed cell wall synthesis to enforce helical curvature (56). Since we showed that halofilins are required to maintain the integrity of the cell envelope during shape-shifting, we hypothesize that HalA and HalB could recognize extreme curvatures in the membrane to prevent cells from collapsing as they bend at fragile cell locales.

Therefore, we revisited our super-resolution microscopy dataset of membrane-stained cells expressing HalA-msfGFP and HalB-msfGFP (Fig. 3A, Movie S2) and extracted sub-micron Gaussian Curvature (GC) measurements mapped across XYZ positions (Fig. 6A). Next, we correlated this curvature profile to the normalized intensity profile map of each halofilin fusion, identifying the curvature regions where there is accumulation of each halofilin at pixel (<300 nm) resolution. Notably, both HalA-msfGFP and HalB-msfGFP showed a strong preference for positive and negative curvatures, respectively (Fig. 6B). Consistent with our previous phenotypic analysis, their accumulation was more pronounced in rods (HalA) or disks (HalB), suggesting that their physical interaction, if it exists, should be transient in a defined cell type. These results support our hypothesis that HalA and HalB could be required for maintaining proper cell curvature balance at critical locales in the membrane.

**Figure 6.**
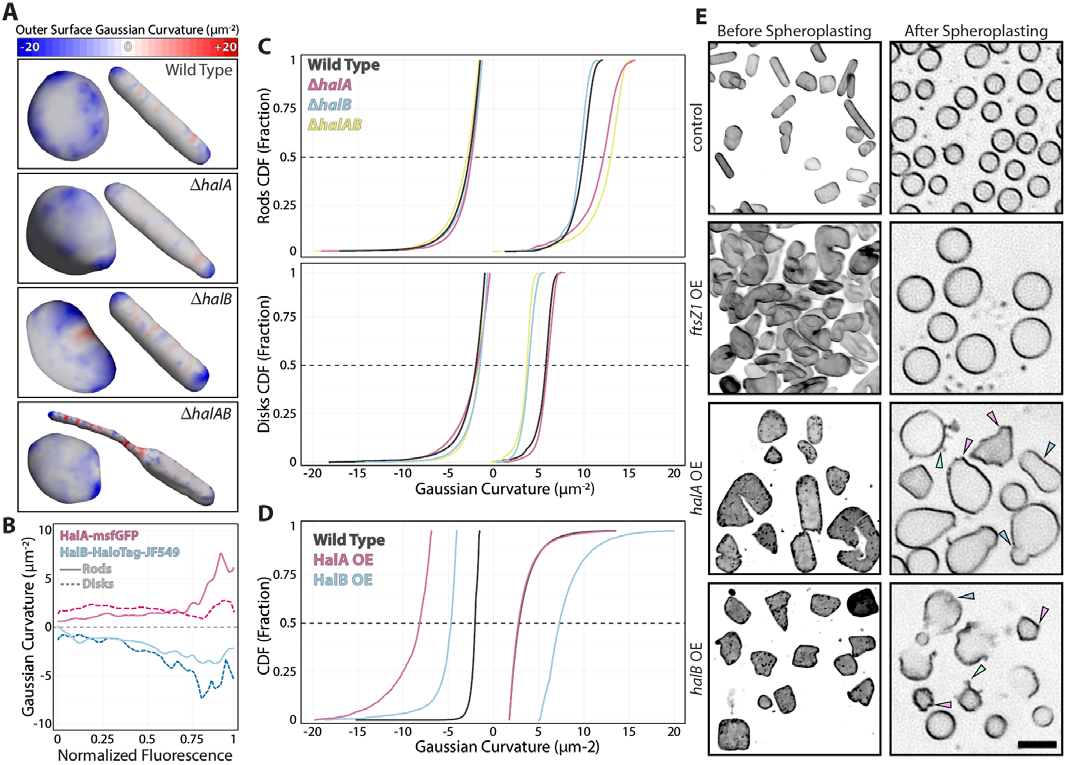
HalA and HalB mediate direct membrane remodeling. **(A)** False-colored, sub-micron mapping across halofilin mutants compared to their parental wild type. Representative cells were arbitrarily chosen, and their membrane curvature scale (from the perspective of its inner surface) was labeled in red (positive), white (neutral), and blue (negative). **(B)** Correlation between spatial localization of membrane curvatures and fluorescence intensity profile shows the affinity of HalA-msfGFP for positive (cytoplasmic surface) and HalB-msfGFP for negative (extracellular surface) membrane curvatures. **(C)** Cumulative distributions of each dataset’s 25% extreme negative and positive Gaussian curvatures. Dashed lines indicate the median values of the distribution. **(D)** Cumulative distributions of cells overexpressing halA and halB under the control of the Pxyl promoter. Cells were grown under induction with 10 mM xylose until reaching mid-exponential phase. **(E)** Spheroplasting wild-type cells and cells overexpressing ftsZ1, halA and halB. in the presence (left column) and absence (right column) of S-layer lattice. Overexpression carried out under 10 mM xylose until mid-exponential. Spheroplasts were obtained by the addition of 50 mM EDTA. 3D-SoRa images were created from maximum intensity projections. Arrowheads indicate angular corners (pink), constricting regions (blue), and inside-out protrusions (green) Scale bar represents 5 µm. Statistical differences were tested by Kolmogorov-Smirnov (P<0.0001).

### Halofilin deletions affect envelope curvature profiles

The enrichment of HalA-msfGFP and HalB-msfGFP at positive and negative GCs, respectively, indicated that Δ*halA* and Δ*halB* morphological defects should correlate with the loss of specific membrane curvatures in rods and disks. To test this hypothesis, we revisited our super-resolution dataset for wild-type and mutant strains (Fig. 2A, Movie S1) to extract submicron membrane curvature maps across different cell types (Fig. 6A). The GC distributions showed a predominance (>50%) of neutral curvatures in both rods and disks. To visualize subtle but crucial differences in our samples, we curated our datasets and selected only the top 25th percentile of both negative and positive curvatures from each dataset.

As a result, we found that the mutant rods showed only slight differences in negative curvatures across Δ*halA*, Δ*halB*, and Δ*halAB* strains, observed only below the 10th percentile (Fig. 6C, top- and bottom-left panels). However, Δ*halA* and Δ*halAB* rods were pronouncedly shifted towards positive curvatures (12.2±1.7 and 13.3±1.9 µm^-2^, respectively) compared to wild type (10±1.2 µm^-2^). In contrast, the positive curvatures of the Δ*halB* rod cells were again alike the wild type (9.5±0.8 µm^-2^) (Fig. 6C, top-right panel). Nevertheless, the disk curvature distributions revealed a recurring pattern of dominance reversal: Δ*halA* cells approximated the wild-type distribution (5.2±0.5 µm^-2^ and 5.5±0.6 µm^-2^, respectively), while Δ*halB* and Δ*halAB* skewed from extreme positive curvatures to flatter surfaces (3.6±0.5 and 3.8±0.9 µm^-2^, respectively) (Fig. 6C, bottom panels). Hence, HalA must be capable of localizing to specific regions to prevent the increase of extreme positive curvatures (above 10 µm^-2^) in rods. In contrast, HalB maintains the balance of negative and positive curvatures, preventing them from becoming flatter. Finally, the transient negative-dominant profile of Δ*halA* over Δ*halAB* rods recapitulates the cell shape measurements shown in Figure 2B, suggesting that poor maintenance of membrane curvature during shape-shifting is responsible for the morphological defects from halofilin mutants.

### Halofilins remodel the membrane independently of the S-layer lattice

Although preferential localization to specific curvatures could facilitate proteins in engaging in membrane remodeling, it is not sufficient by itself to conclude that halofilins are actively involved in creating specific curvatures *in vivo*. To test if halofilins could individually deform cells, we drove the overexpression of *halA* and *halB* using the strong and inducible promoter Pxyl (57). In this experiment, we expected to observe cell curvature distributions in direct opposition to the ones recorded from halofilin mutants; i.e., an excess of HalA would shift Gaussian curvatures away from extreme positives, and high levels of HalB would push membrane deformation towards extreme curvatures. Cultures overexpressing *halA* (*halA* OE) and *halB* (*halB* OE) generated mostly larger, deformed cells compared to wild type (5.7±1.4 µm^-2^, 5.6±1.5 µm^-2^, and 2.6±0.8 µm^-2^, respectively) (Figs. 6D and S13A-B, Movie S1). From 3D projections, we calculated Gaussian curvatures and observed an increase of negative curvatures from *halA* OE compared to wild type (−8.2±3.3 µm^-2^ and - 2.0±1.1 µm^-2^, respectively), but no differences in positive curvatures (2.8±0.9 µm^-2^ and 2.7±0.8 µm^-2^, respectively). Nonetheless, *halB* OE induced shifts in both extreme negative (−4.3±1.7 µm^-2^) and positive (7.2±3.5 µm^-2^) directions, supporting the hypothesis that HalB acts on maintaining the curvature balance away from flatter surfaces.

Even though the absence and excess of halofilins did not falsify our mechanical scaffolding model, the data above do not rule out the recruitment of enzymes acting directly or indirectly in the S-layer remodeling. To differentiate between these two ideas, we asked whether deformed cells overexpressing halA and halB could preserve membrane curvatures independently of the S-layer lattice. To do this, we again removed the S-layer by spheroplasting as in Fig. 3F. As controls, we compared both wild-type cells and a strain overexpressing ftsZ1 (ftsZ1 OE). ftsZ1 OE is a convenient control as its overexpression results in deformed cells similarly to halA OE and halB OE (18, 57). Moreover, FtsZ proteins have cytomotive functions, and overexpression of ftsZ in bacteria can deform spheroplasts only under iso-osmotic conditions (58). Thirty minutes after the addition of EDTA to Hfx. volcanii to form spheroplasts, we observed that wild-type cells exhibited uniformly spherical morphology (Fig. 6E). Interestingly, ftsZ1 OE cells also resulted in regular spheres, but larger sizes compared to the wild type, as observed before spheroplasting. However, in halA OE and halB OE backgrounds, the spheroplasts showed irregular spheroids with lower solidity (Fig. S13C). We also observed a diversity of recurring membrane patterns, such as angular corners, constricting regions, and inside-out protrusions. Strikingly, while halA OE and halB OE spheroplast curvatures were qualitatively comparable, halB OE cells showed sub-populations of broken cells. This observation aligns with our curvature measurements from halB OE cells, where cells accumulated a diversity of both extreme negative and positive curvatures, which could result in membrane breakage and subsequent lysis (Fig. 6D).

## Discussion

Walled cells have evolved the ability to confine higher intracellular pressure, albeit at the cost of having to slowly build *de novo* cell wall during morphological recovery or development over one or more generations (59, 60). By probing the function of the halofilins HalA and HalB, we have drawn a parallel to canonical bactofilins, such as a preference for specific membrane curvatures to prevent cells from warping at critical regions (Fig. 6B). However, disk-to-rod shape transitions in *Hfx. volcanii* occur within a quarter of its generation time (Fig. 5F). This argument, combined with spheroplast experiments showing that halofilins alone are sufficient to produce membrane deformation (Fig. 6E), supports the model where halofilins act directly at the envelope as mechanical polymers. Nevertheless, it is not possible at this point to rule out the possibility of a hybrid mechanism between the recruitment of envelope enzymes and mechanical scaffolding.

The observation that the deletion of halofilins result in decondensation of the Z-ring at the midcell leaves open questions for future research (Fig. 5A-B). Although the resulting defects are likely due to membrane curvature aberrations created during shape transitions, we can speculate that the spatiotemporal control of negative and positive curvatures across the cell could be an alternative path of positioning the cell division machinery across different cell shapes. Supporting this idea, we also observed that HalA-msfGFP accumulates preferentially around the midcell (Fig. 3B). The idea of controlled membrane deformation playing a role in Z-ring positioning aligns with the absence of a dedicated Min system in archaea (61) and mathematical models linking cell shape to Turing patterns (62). Moreover, this could expand our understanding of Z-ring positioning universally across bacteria, where cells direct cell division placement with high accuracy even when both Min and nucleoid occlusion systems are deleted (63–65).

A central outstanding question in the field is how closely related haloarchaeal species build different cell shapes (14). In the past, mutations disrupting the covalent attachment of the SLG to phospholipids induced rod formation, delaying the transition to disks until the very late exponential phase (9). Following this logic, we speculate that the S-layer may have a lower degree of lipidation in rods than in disks, making rods more vulnerable to membrane deformations, which would explain the higher proportion of negative and positive curvatures in rods (Fig. 6C). Disks, on the other hand, are flatter, possibly due to reduced membrane fluidity caused by the attachment to the S-layer lattice. This reasoning reconciles the specific phenotypes from Δ*halA* and Δ*halB* to specific cell types (rods and disks, respectively), which requires further research to substantiate.

Furthermore, little is known about the dynamics of the S-layer and how many SLG lattice conformations exist in rods and disks and during shapeshift between them (66). von Kügelgen and colleagues recently combined cryo-EM and *in situ* cryo-ET to show that the *Hfx. volcanii* S-layer comprises SLG ensembles of symmetric P6 groups but with P5 groups in high curvature regions (47). Interestingly, the bHalB subdomain may be at the same plane as the S-layer, which could explain its static localization compared to HalA in live-cell time-lapses (Fig. 3C, Movie S2). Consequently, bHalB interaction with the S-layer could be mediating lattice reorganization and rigidity in disks on the verge of shapeshift.

In summary, we propose a model of how halofilins contribute to disk-to-rod shape transitions at different, sequential stages (Fig. 7). At the beginning of the shape transition process, HalB could induce lattice remodeling at negative membrane curvatures to counterbalance the rigidity of the membrane attached to the S-layer lattice. Alternatively, HalB could preferentially interact with SLG at disrupted lattice locales, “patching up” the wounds while the envelope is remodeled. Another non-mutually exclusive possibility is that HalB directs the removal of lipidated SLG, promoting the turnover to the unlipidated lattice mostly present in rods. Thereafter, as the shifting cells decrease the levels of lipidation, HalA filaments dynamically accumulate at positive curvatures to prevent membrane Gaussian curvatures above 10µm^-2^. Therefore, when Δ*halB* disk cells fail to control S-layer remodeling in the first stage of development, HalA could partially repair cells by reinforcing positive curvatures. This might explain why we did not observe significant defects in Δ*halB* rods and Δ*halA* disks, the latter possibly connected to rod-to-disk transitions. However, this model requires the characterization of lipidation profiles in different cell types. Future studies should also clarify the details of a possible transient interaction between HalA and HalB during shape transitions, and whether halofilin filaments participate in the detection of (currently unknown) environmental cues triggering shape development. Identification of new factors and structural studies on S-layer lattices should play a pivotal role in this process. Ultimately, further mechanistic details of how archaeal cytoskeleton couples mechanosensing to the cell envelope will expand our understanding of the switch from mono-to multi-functional polymers after eukaryogenesis.

**Figure 7.**
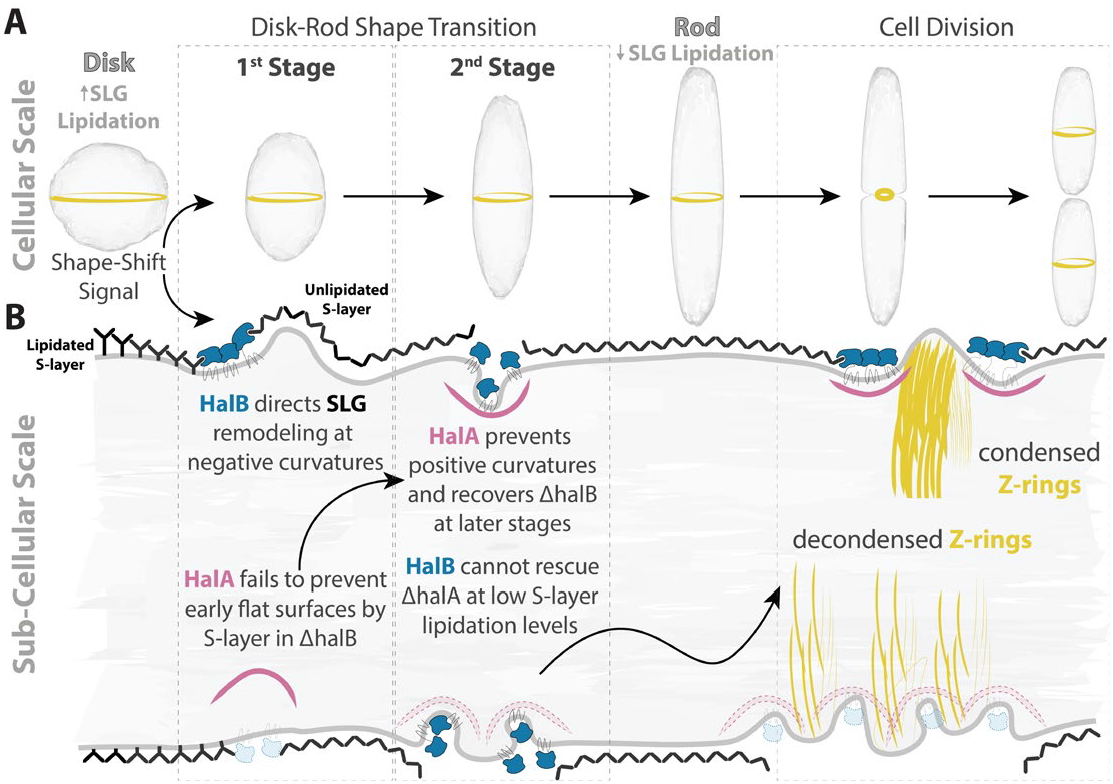
Proposed model for how HalA and HalB act at different time points during disk-rod shape transitions. **(A)** Summary of shape transition stages at the cellular scale. Disks sense the (still elusive) signal from the environment and trigger the shape-shift developmental program toward rod cells. Cell division resumes after the shape transition is finalized. The division site (Z-ring) is represented in yellow. **(B)** Zoom-in of the proposed molecular mechanism of HalA (pink) and HalB (blue) roles during shape transition at a sub-cellular scale. The dark grey outline represents the cytoplasmic membrane and the S-layer lattice altogether. The upper envelope layer represents the process in wild-type cells, while the bottom flow illustrates the scenarios in the ΔhalA and ΔhalB strains. Z-ring yellow threads represent FtsZ1 and FtsZ2 protofilaments. A speculative HalA:HalB interaction, if existent at all, should be transient during development. The proposed lipidation role in the model is still experimentally untested.

## Materials and Methods

A detailed description of protocols, and reagents, as well data and software availability can be found in Supporting Information.

## Acknowledgments

The authors thank Bruce Goode and Avital Rodal (Brandeis U) for their comments on our manuscript. The Bisson Lab appreciates the Goode Lab (Brandeis U) for access to their SoRa microscope and the sharing of their pMAL vector. Revisions on this work in a timely matter were only possible with the support of the Brandeis Light Microscopy Core Facility and the Louise Mashal Gabbay Cellular Visualization Center at Brandeis U. OL is grateful to Benjamin Bratton (Vanderbilt U) and Thomas Fai (Brandeis U) for their advice on Gaussian Curvature analysis. AB would like to acknowledge Jan Löwe prompt advice in obtained and TEM imaging of pure fractions of bactofilins. JM would also like to thank TJ Koehler for his help with navigating the Blender software. This work was supported by the Moore–Simons Project on the Origin of the Eukaryotic Cell (doi:10.46714/735929LPI) awarded to AB; the Human Frontiers Science Program (RGY0074/2021) awarded to AB and VA; the National Science Foundation Grant (NSF-MBC2222076) awarded to MP and AB; and the Brandeis National Science Foundation (NSF) Materials Research Science and Engineering Center (MRSEC) Bioinspired Soft Materials (NSF-DMR 2011846); and NSF-MCB1651117 to AKS. AB is a Pew Scholar in the Biomedical Sciences, supported by The Pew Charitable Trusts. This work was partly supported by institutional funds of the Max Planck Society. VA would like to thank Andrei Lupas (MPI Tübingen) for their continued support.

