## Supplemental Materials for "Halofilins as Emerging Bactofilin Families of Archaeal Cell Shape Plasticity Orchestrators"

PNAS

**This PDF file includes:**

- Material and Methods
- Figures S1 to S13
- Tables S1 to S5
- Legends for Movies S1 to S6
- SI References

**Other supporting materials for this manuscript include the following:**

- Movies S1 to S6

### Supporting Information Text

### Material and Methods

#### Homo-oligomer structural prediction

We used an installation of AlphaFold2-Multimer (v.2.3.1) (1, 2) at the Max Planck Computing and Data Facility to create homo-oligomer structure predictions. We computed dimer, trimer, and tetramer predictions for TtBac (UniProtKB Q72HS6); dimer, trimer, and tetramer predictions for HalA (UniProtKB D4GZ39/HVO_1610); dimer, trimer, and tetramer predictions for bHalB (UniProtKB D4GX31/HVO_1237, residues 41-125); and dimer, trimer, tetramer, pentamer, and hexamer predictions for the TM domain of HalB (UniProtKB D4GX31/HVO_1237, residues 152-297). All predictions were carried out using the default settings.

#### Structure alignments and multiple sequence alignments

We performed protein (multiple) structure alignments using US-align (3) with default settings. The structure-guided MSA shown in Figure S3, was retrieved from the output of US-align, manually curated, and visualized with pyMSAviz (https://github.com/moshi4/pyMSAviz).

#### Alignment of profile HMMs

An HHpred search was run for HalA (UniProtKB D4GZ39/HVO_1610) against the Pfam-A (v.36) (4) profile HMM database. Individual HHpred alignments were conducted between bHalB (UniProtKB D4GX31/HVO_1237, residues 41-125) and TtBac (UniProtKB Q72HS6) and between bHalB and HalA. All HHpred jobs were run using the MPI Bioinformatics Toolkit (5). The secondary structure scoring parameter was disabled to rule out the chance that incidental similarities influenced high scores for matches in their secondary structures. Profile HMMs were built for each sequence by running three iterations of HHblits (6) against the Uniclust30 database (7), using an E-value cutoff set to 1e-3.

#### Cluster map generation

The cluster map shown in Figure 1A was created by first performing a local BLASTp (v.2.13.0+) (8) search against the UniProtKB (release 2023_04, of 13-Sep-2023) (9). We used HalA (UniProtKB D4GZ39/HVO_1610) and HalB (UniProtKB D4GX31/HVO_1237) as queries and an E-value cutoff set to 10. The protein sequences found in the previous search were used as seeds for a second iteration of BLASTp against UniProtKB with the same E-value cutoff to gather more distant homologs of our initial queries. We inspected the results manually and browsed the predicted structure of an arbitrary number of sequence hits at the UniProtKB web server using their respective accessions as queries. We chose 8 final representative seeds per halofilin, which allowed us to capture the structural and taxonomic diversity within each respective family. We further collected 18 protein sequences of previously reported canonical bactofilins and queried the UniProtKB web server for entries containing the term “bactofilin,” retrieving the 2 matches issued from Swiss-Prot (Table S1). To gather more sequences for cluster analysis, we performed a local BLASTp search against the UniProtKB, using these 36 seed sequences as queries and an E-value cutoff set to 10. We only kept the BLASTp hits whose E-value was ≤ 1e-3 and query coverage per subject was ≥ 50%. Duplicates, fragments, and partial sequences were removed. All sequences that passed the filtering criteria were pooled together, totaling 8,542 protein sequences. Pairwise sequence similarities were computed for all protein sequences using BLASTp, with an E-value cutoff set to 1. Clustering was achieved with CLANS (10), using the pairwise BLASTp E-values calculated in the previous step. CLANS ran for 100,000 iterations, achieving convergence, with an E-value cutoff of 1e-10 and a repulse value set to 60. The edges in the map are shown for E-values ≤ 1e-8. Data analysis was carried out with Pandas v.1.5.3 (https://github.com/pandas-dev/pandas), Biopython v.1.81 (https://github.com/biopython/biopython), and NetworkX v.3.0 (https://github.com/networkx/networkx), and the cluster map was plotted using Matplotlib v.3.7.1 (https://github.com/matplotlib/matplotlib). These packages were installed and run under Python v.3.10.12 (https://www.python.org/). Structure visualizations were created in PyMOL v.2.5.0 (https://github.com/schrodinger/pymol-open-source).

#### Structural modeling of HalB

The SignalP-6.0 (11) web server (https://services.healthtech.dtu.dk/services/SignalP-6.0/) was used to predict the presence of a signal peptide in HalB (UniProtKB D4GX31/HVO_1237). SignalP-6.0 detected a signal peptide (Pr = 0.9778) with a cleavage site between residues 29-30. Using this information, the structural model of HalB lacking the signal peptide (residues 31-321) was built using an installation of AlphaFold2 (v.2.3.1) (1) at the Max Planck Computing and Data Facility. The prediction was carried out using the default settings, and the model with the highest confidence (ranked_0.pdb) was chosen for downstream use. The MembraneFold web server (https://ku.biolib.com/MembraneFold) (12) was used to predict the membrane positioning of HalB, using the PDB file for the structural model generated by AlphaFold2 as input. Structure visualizations were created in PyMOL v.2.5.0.

#### Taxonomy

We parsed the FASTA header of each of the 8,542 sequences in our dataset, and in case the sequence was issued from UniProtKB, we retrieved the respective NCBI TaxId from the "OX" field. If a sequence was issued from NCBI's protein sequence database (13), we retrieved the organism's scientific name instead. We then converted it to its respective TaxId using the "name2taxid" subcommand from TaxonKit (v.0.14.1) (14). Afterward, we used TaxonKit's "reformat" subcommand to retrieve the taxonomic lineages in canonical ranks, given these TaxIds as input, with the "--miss-rank-repl" flag set to "unclassified". The NCBI Taxonomy database dump files used in conjunction with TaxonKit were downloaded on 18-Oct-2023 from https://ftp.ncbi.nih.gov/pub/taxonomy/taxdump.tar.gz. Taxonomy data was analyzed with Pandas v.1.5.3, and plots were generated with Plotly v.5.17.0 (https://github.com/plotly/plotly.py) under Python v.3.10.12.

#### Media and Culture Growth

*Hfx. volcanii* strains were grown in the semi-defined Hv-Cab medium (15). Strains were streaked from -80°C freezer stocks onto Hv-Cab plates (1.5% agar), supplemented with 20 µM tryptophan and 50 µM uracil when necessary, and incubated for 2-3 days at 45°C. Single colonies were transferred into 16×25 mm glass tubes with 3 mL of Hv-Cab, placed on a roller drum, and grown at 42°C to mid-exponential phase ($OD_{600nm}$~0.5), resulting in a mixed population of rod- and disk-shaped cells. Cultures were grown to early-exponential (rods, OD600nm~0.2) or late-exponential (disks, OD600nm~0.8) phases when specific cell shapes were required. The expression of HalA- and HalB fluorescent fusions was induced by adding 500 µM Tryptophan. Overexpression of HalA and HalB was carried out under 5mM xylose (16). *Hbt. salinarum* strains were grown as described for *Hfx. volcanii*, using the CM media (250 g/L NaCl; 20 g/liter $MgSO_4 \cdot 7H_2O$; 3 g/L trisodium citrate; 2 g/L KCl; 10 g/L Oxoid bacteriological peptone, pH 6.8) supplemented with 10 µg/ml mevinolin in plates and liquid cultures.

#### Cloning and Transformation

Oligos, plasmids and strains used in this study can be found in Tables S3-S5. Plasmids were cloned using isothermal (Gibson) reactions (17) and transformed into competent *E. coli* DH5α cells, transformants selected on LB plates supplemented with 100µg/ml Carbenicillin (RPI Research Products, #4800-94-6). Individual clones were confirmed by whole-plasmid sequencing (Plasmidsaurus, plasmidsaurus.com). Plasmid preps were then transformed into *Hfx. volcanii* using the method previously described (18), using 0.5 M EDTA (Thermo Scientific, #J15694-AE) and PEG600 (Sigma, # 87333-250G-F). *Hfx. volcanii* strains were built in the parental H26 (*ΔpyrE2*) or H53 (*ΔpyrE2 ΔtrpA*) wild-type strains with constructs cloned into the replicative pTA962 and pAL750 vectors (Table S3). Synthetic genes used to amplify fragments to introduce tags, fluorescent proteins, and flexible linkers of different lengths were already previously described, including complete DNA and amino acid sequences (19). Knockout strains were created by the "pop-in, pop-out" strategy (18) using 5-Fluoroorotic Acid (Fisher, #FERR0811) as a counter-selection for *pyrE2* pop-out. A detailed description of each strain used in this work follows:

*aJK3 [ΔpyrE2 pTA962::halA-40aa-msfGFP]* was created by transforming the H26 strain with the eBL94 plasmid, cloned from a Gibson assembly consisting of 3 parts: 1) *halA fragment:* PCR with primers oBL182 and oTR02 from H26 gDNA template; 2) *40aa-msfGFP fragment*: PCR with primers oBL354 and oBL31 from a synthetic DNA template; 3) pTA962 plasmid linearized with NdeI.

*aBL404 [ΔpyrE2 pTA962::halB-40aa-msfGFP]* was created by transforming H26 with the eBL301 plasmid, cloned from a Gibson assembly consisting of 3 parts: 1) *halB fragment:* PCR with primers oBL397 and oBL198 from H26 gDNA template; 2) *40aa-msfGFP fragment*: PCR with primers oBL354 and oBL31 from a synthetic DNA template; 3) pTA962 plasmid linearized with NdeI.

*aBL369 [ΔpyrE2 pTA962::halB-30aa-HaloTag(Ct)]* was created by transforming H26 with the eBL271 plasmid, cloned from a Gibson assembly consisting of 3 parts: 1) *halB fragment:* PCR with primers oBL397 and oBL198 from H26 gDNA template; 2) *30aa-HaloTag fragment*: PCR with primers oBL354 and oBL318 from a synthetic DNA template; 3) pTA962 plasmid linearized with NdeI.

*aBL370 [ΔpyrE2 pTA962::halB-30aa-HaloTag(SW)]* was created by transforming H26 with the eBL272 plasmid, cloned from a Gibson assembly consisting of 4 parts: 1) *halB fragment A:* PCR with primers oBL397 and oHV186 from H26 gDNA template; 2) *15aa-HaloTag-15aa fragment*: PCR with primers oHV84 and oHV82 from a synthetic DNA template; 3) *halB fragment B:* PCR with primers oHV188 and oBL472 from H26 gDNA template; 4) pTA962 plasmid linearized with NdeI.

*aKA11 [ΔpyrE2 pTA962::HaloTag]* was created by transforming H26 with the eKA3 plasmid, cloned from a Gibson assembly consisting of 2 parts: 1) *HaloTag fragment:* PCR with primers oHV36 and oBL318 from a synthetic DNA template; 3) pTA962 plasmid linearized with NdeI.

*aBL439 [ΔpyrE2 pTA962::halA-30aa-HaloTag]* was created by transforming H26 with the eBL319 plasmid, cloned from a Gibson assembly consisting of 3 parts: 1) *halA fragment:* PCR with primers oBL397 and oBL198 from H26 gDNA template; 2) *30aa-HaloTag fragment*: PCR with primers oBL354 and oBL318 from a synthetic DNA template; 3) pTA962 plasmid linearized with NdeI.

*aBL336 [ΔpyrE2 pTA962::halA-30aa-HaloTag pTA962::halA-ΔMTS-40aa-msfGFP]* was created by co-transforming H26 with the eBL319 plasmid described above together with eBL232 plasmid, cloned from a Gibson assembly consisting of 5 parts: 1) *halA fragment A:* PCR with primers oBL397 and oBL359 from H26 gDNA template; 2) *40aa fragment*: synthetic fragment; 3) *halA fragment B:* PCR with primers oBL360 and oHV189 from H26 gDNA template; 4) *40aa-msfGFP fragment*: PCR with primers oBL354 and oBL31 from a synthetic DNA template; 5) pTA962 plasmid linearized with NdeI.

*aBL416 [ΔpyrE2 pTA962::halA-30aa-HaloTag pTA962::msfGFP-MTS-mYPet]* was created by co-transforming H26 with the eBL319 plasmid described above together with eBL311 plasmid, cloned from a Gibson assembly consisting of 3 parts: 1) *msfGFP-MTS-mYPet:* synthetic fragment; 2) *40aa-msfGFP fragment*: PCR with primers oBL354 and oBL31 from a synthetic DNA template; 3) pTA962 plasmid linearized with NdeI.

*aBL420 [ΔpyrE2 pTA962::halA-30aa-HaloTag pTA962::halA(Nt)-40aa-msfGFP]* was created by co-transforming H26 with the eBL319 plasmid described above together with eBL313 plasmid, cloned from a Gibson assembly consisting of 3 parts: 1) *halA(Nt) fragment:* PCR with primers oBL397 and oBL433 from H26 gDNA template; 2) *40aa-msfGFP fragment*: PCR with primers oBL354 and oBL31 from a synthetic DNA template; 3) pTA962 plasmid linearized with NdeI.

*aBL423 [ΔpyrE2 pTA962::halA-30aa-HaloTag pTA962::halA(Nt)(ΔMTS)-40aa-msfGFP]* was created by co-transforming H26 with the eBL319 plasmid described above together with eBL314 plasmid, cloned from a Gibson assembly consisting of 3 parts: 1) *halA(Nt)(ΔMTS) fragment:* PCR with primers oBL397 and oBL357 from H26 gDNA template; 2) *40aa-msfGFP fragment*: PCR with primers oBL354 and oBL31 from a synthetic DNA template; 3) pTA962 plasmid linearized with NdeI.

*aBL401 [ΔpyrE2 pTA962::halA-30aa-HaloTag pTA962::halA(Ct)-40aa-msfGFP]* was created by co-transforming H26 with the eBL319 plasmid described above together with eBL299 plasmid, cloned from a Gibson assembly consisting of 2 parts: 1) *halA(Ct) fragment:* PCR with primers oBL426 and oHV189 from H26 gDNA template; 2) pTA962 plasmid linearized with NdeI.

*aBL412 [ΔpyrE2 pAL750::halA]* was created by transforming H26 with the eBL307 plasmid, cloned from a Gibson assembly consisting of 2 parts: 1) *halA fragment:* PCR with primers oBL436 and oBL440 from H26 gDNA template; 2) pAL750 plasmid linearized with NdeI.

*aBL414 [ΔpyrE2 pAL750::halB]* was created by transforming H26 with the eBL309 plasmid, cloned from a Gibson assembly consisting of 2 parts: 1) *halB fragment:* PCR with primers oBL441 and oBL442 from H26 gDNA template; 2) pAL750 plasmid linearized with NdeI.

*aBL80 [Δtrp ΔpyrE2 ΔhalA]* was created by transforming H53 with the pTA131::halA suicide plasmid, cloned by DNA Ligase reaction consisting of 3 parts: 1) halA upstream fragment: PCR with primers FW_up_halA and RV_up_halA from H53 gDNA template; 2) PCR with primer FW_down_halA and RV_down_halA from H53 gDNA template; 3) pTA131 plasmid linearized with the HindIII and XbaI.

*aZC85 [Δtrp ΔpyrE2 ΔhalB]* was created by transforming H53 with the eZC69 suicide plasmid, cloned from a Gibson assembly consisting of 4 fragments: 1) *halB* upstream fragment: PCR with primers oBL369 and oZC44 from DS2 genomic DNA template; 2) *halB* downstream fragment: PCR with primers oZC40 and oZC41 from DS2 genomic DNA template; 3) *pyrE2* cassette fragment: PCR with primers oZC42 and oZC43 from pTA962 plasmid DNA template; 4) ampR-ori_f1 fragment: PCR with primers oBL105 and oBL97 from pBluescript II (Agilent) plasmid DNA template. Transformants were streaked on Hv-Cab plates incubated at 45℃. Colonies were grown in liquid Hv-Cab overnight, back diluted into Hv-Cab, and grown overnight. This step was repeated, and cultures were back diluted into HV-Cab supplemented with 50 µM uracil, grown overnight, then back diluted and regrown in Hv-Cab plates supplemented with uracil. Cultures were streaked on Hv-Cab plates supplemented with 50µM uracil and 50µg/ml 5-FOA. Survivors were confirmed for *halB* deletion by PCR.

*aZC37 [Δtrp ΔpyrE2 ΔhalA ΔhalB]* was created by transforming aBL80 with the eZC69 plasmid and repeating the process described above.

*aBL327 [ΔhalA ΔpyrE2 ΔtrpA pTA962:: halA-msfGFP]* was created by transforming aBL80 with eBL94 plasmid described above.

*aZC41 [ΔHalB pTA962::halB-40aa-msfGFP]* was created by transforming aZC36 with eBL301 plasmid described above.

*aZC65 [Δtrp ΔpyrE2 ΔhalA ΔpyrE2 pTA962::ftsZ1-GFP]* was created by transforming aBL80 with pIDJL40, a gift from Iain Duggin.

*aZC67 [ΔpyrE2 ΔhalB ΔpyrE2 pTA962::ftsZ1-GFP]* was created by transforming aZC36 with pIDJL40, a gift from Iain Duggin.

*aZC69 [Δtrp ΔpyrE2 ΔhalA ΔhalB ΔpyrE2 pTA962::ftsZ1-GFP]* was created by transforming aZC37 with pIDJL40, a gift from Iain Duggin.

*aZC71 [Δtrp ΔpyrE2 ΔhalA ΔpyrE2 pTA962::ftsZ2-GFP]* was created by transforming aBL80 with pHVID95, a gift from Iain Duggin.

*aZC73 [ΔpyrE2 ΔhalB ΔpyrE2 pTA962::ftsZ2-GFP]* was created by transforming aZC36 with pHVID95, a gift from Iain Duggin.

*aZC75 [Δtrp ΔpyrE2 ΔhalA ΔhalB ΔpyrE2 pTA962::ftsZ2-GFP]* was created by transforming aZC37 with pHVID95, a gift from Iain Duggin.

*aZC99 [ΔcetZ1 ΔtrpA ΔpyrE2 pTA962::halA-40aa-msfGFP]* was created by transforming aBL456 (Δ*cetZ1* Δ*trpA* Δ*pyrE2*, a gift from Thane Papke) with the eBL94 plasmid described above.

*aZC101 [ΔcetZ2 ΔtrpA ΔpyrE2 pTA962::halA-40aa-msfGFP]* was created by transforming aBL456 (Δ*cetZ2* Δ*trpA* Δ*pyrE2*, a gift from Thane Papke) with the eBL94 plasmid described above.

*aZC103 [ΔcetZ3 ΔtrpA ΔpyrE2 pTA962::halA-40aa-msfGFP]* was created by transforming aBL460 (Δ*cetZ3* Δ*trpA* Δ*pyrE2*, a gift from Thane Papke) with the eBL94 plasmid described above.

*aZC105 [ΔcetZ4 ΔtrpA ΔpyrE2 pTA962::halA-40aa-msfGFP]* was created by transforming aBL462 (Δ*cetZ4* Δ*trpA* Δ*pyrE2*, a gift from Thane Papke) with the eBL94 plasmid described above.

*aZC107 [ΔcetZ5 ΔtrpA ΔpyrE2 pTA962::halA-40aa-msfGFP]* was created by transforming aBL464 (Δ*cetZ5* Δ*trpA* Δ*pyrE2*, a gift from Thane Papke) with the eBL94 plasmid described above.

*aZC109 [ΔcetZ6 ΔtrpA ΔpyrE2 pTA962::halA-40aa-msfGFP]* was created by transforming aBL454 (Δ*cetZ6* Δ*trpA* Δ*pyrE2*, a gift from Thane Papke) with the eBL94 plasmid described above.

*aZC111 [ΔvolA ΔpyrE2 pTA962::halA-40aa-msfGFP]* was created by transforming aKA16 (Δ*volA* Δ*pyrE2)* with the eBL94 plasmid described above.

*aZC115 [ΔftsZ1 ΔftsZ2 ΔhdrB ΔpyrE2 pTA962::halA-40aa-msfGFP]* was created by transforming ID112, (Δ*ftsZ1* Δ*ftsZ2* Δ*hdrB* Δ*pyrE2,* a gift from Iain Duggin), with the eBL94 plasmid described above.

*aZC87 [ΔcetZ1 ΔtrpA ΔpyrE2 pTA962::halB-40aa-msfGFP]* was created by transforming aBL456 (Δ*cetZ1* Δ*trpA* Δ*pyrE2*, a gift from Thane Papke) with the eBL301 plasmid described above.

*aZC89 [ΔcetZ2 ΔtrpA ΔpyrE2 pTA962::halB-40aa-msfGFP]* was created by transforming aBL458 (Δ*cetZ2* Δ*trpA* Δ*pyrE2*, a gift from Thane Papke) with the eBL301 plasmid described above.

*aZC91 [ΔcetZ3 ΔtrpA ΔpyrE2 pTA962::halB-40aa-msfGFP]* was created by transforming aBL460 (Δ*cetZ3* Δ*trpA* Δ*pyrE2*, a gift from Thane Papke) with the eBL301 plasmid described above.

*aZC93 [ΔcetZ4 ΔtrpA ΔpyrE2 pTA962::halB-40aa-msfGFP]* was created by transforming aBL462 (Δ*cetZ4* Δ*trpA* Δ*pyrE2*, a gift from Thane Papke) with the eBL301 plasmid described above.

*aZC95 [ΔcetZ5 ΔtrpA ΔpyrE2 pTA962::halB-40aa-msfGFP]* was created by transforming aBL464 (Δ*cetZ5* Δ*trpA* Δ*pyrE2*, a gift from Thane Papke) with the eBL301 plasmid described above.

*aZC97 [ΔcetZ6 ΔtrpA ΔpyrE2 pTA962::halB-40aa-msfGFP]* was created by transforming aBL454 (Δ*cetZ6* Δ*trpA* Δ*pyrE2*, a gift from Thane Papke) with the eBL301 plasmid described above.

*aZC113 [ΔvolA ΔpyrE2 pTA962::halB-40aa-msfGFP]* was created by transforming aKA16 (Δ*volA* Δ*pyrE2)* with the eBL301 plasmid described above.

*aZC117 [ΔftsZ1 ΔftsZ2 ΔhdrB ΔpyrE2 pTA962::halB-40aa-msfGFP]* was created by transforming ID112, (Δ*ftsZ1* Δ*ftsZ2* Δ*hdrB* Δ*pyrE2,* a gift from Iain Duggin), with the eBL301 plasmid described above.

*aKA16 [ΔpyrE2 ΔvolA]* was created by transforming H26 with the eTR64 suicide plasmid, cloned from a Gibson assembly consisting of 3 fragments: 1) *volA* upstream fragment: PCR with primers oTR202 and oTR203 from DS2 genomic DNA template; 2) *volA* downstream fragment: PCR with primers oTR204 and oTR205 from DS2 genomic DNA template; 3) *pyrE2*_ampR-ori_f1 fragment: PCR with primers oTR201 and oBL97 from eTR58 (derived from pBluescript II (Agilent)) plasmid DNA template. Cells were plated and transformants struck onto Hv-Cab plates incubated at 45°C. Colonies were grown in liquid Hv-Cab overnight, back diluted into Hv-Cab, and grown overnight. Cultures were then back diluted into Hv-Cab supplemented with 50 μM uracil, grown overnight, then back diluted again in Hv-Cab supplemented with 50 μM uracil. Cultures were struck onto Hv-Cab plates supplemented with 50 μM uracil and 50 μg/ml 5-FOA. Surviving colonies were confirmed for volA deletion by PCR. After positive confirmation of one colony (stocked as aKA16), volA deletion was confirmed by Illumina.

*aBL469 [Δura3 ΔhalA]* was created by transforming the Δura3 strain with the eBL330 suicide plasmid, cloned from a Gibson assembly consisting of 3 fragments: 1) *halA* upstream fragment: PCR with primers oBL477 and oBL478 from NRC-1 genomic DNA template; 2) *halA* downstream fragment: PCR with primers oBL479 and oBL480 from NRC-1 genomic DNA template; 3) pNBKO7 vector plasmid linearized with EcoRI. Transformation was carried out using the standard double-crossover counterselection method previously described (20) and potential transformed strains were selected first, on CM plates without uracil addition, but containing mevinolin (10 μg/ml). The resulting merodiploid strains were then plated on CM plates containing 5-fluoroorotic acid (250 μg/ml) and uracil (50 mg/l) to remove the integrated plasmid, yielding unmarked Δ*halA* deletion strains.

*aBL556 [Δura3 pMTCHA]* was created by transforming Δ*ura3* strain with the pMTCHA plasmid and selected on CM plates supplemented with 10 μg/ml mevinolin and uracil.

*aBL558 [Δura3 ΔhalA pMTCHA]* was created by transforming aBL469 with the pMTCHA plasmid and selected on CM plates supplemented with 10 μg/ml mevinolin and uracil.

*aBL560 [Δura3 pMTCHA::halA]* was created by transforming Δ*ura3* strain with the eZC98 plasmid, cloned from a Gibson assembly consisting of 2 fragments: 1) *halA* gene: PCR with primers oBL481 and oBL482 from NRC-1 genomic DNA template; 2) pMTCHA vector linearized with NdeI, and selected on HCM plates supplemented with 10 μg/ml mevinolin and uracil.

*aBL562 [Δura3 ΔhalA pMTCHA::halA]* was created by transforming aBL469 with the eZC98 plasmid described above, and selected on HCM plates supplemented with 10 μg/ml mevinolin and uracil.

*aJM140 [ΔpyrE2 pyrE2::P250-msfGFP-mevR]* was created by transforming H26 with the eJM53 suicide plasmid, cloned from a Gibson assembly consisting of 5 fragments: 1) *pyrE2* upstream fragment: PCR with primers oBL136 and oBL87 from DS2 genomic DNA template; 2) *pyrE2* downstream fragment: PCR with primers oHV67 and oBL137 from DS2 genomic DNA template; 3) mevinolin resistance fragment: PCR with primers oBL355 and oJM160 from pMTCHA DNA template; 4) P250-msfGFP fragment: PCR with primers oJM212 and oJM213 from eBL227 template 5) ampR-ori_f1 fragment: PCR with primers oBL105 and oBL97 from pBluescript II (Agilent) plasmid DNA template. Transformants were selected on Hv-Cab plates supplemented with 10ug/ml mevinolin and uracil.

*aBL564 [ΔpyrE2 pyrE2::P250-msfGFP-mevR pTA962]* was created by transforming aJM140 with the pTA962 plasmid.

*aBL565 [ΔpyrE2 pyrE2::P250-msfGFP-mevR pTA962::halA-40aa-msfGFP]* was created by transforming aJM140 with the eBL94 plasmid described above.

*aBL566 [ΔpyrE2 pyrE2::P250-msfGFP-mevR pTA962::halA(Nt)(ΔMTS)-40aa-msfGFP]* was created by transforming aJM140 with the eBL319 plasmid described above.

*aBL567 [ΔpyrE2 pyrE2::P250-msfGFP-mevR pTA962::halB-40aa-msfGFP]* was created by transforming aJM140 with the eBL301 plasmid described above.

*eJM71 [BL21(DE3) pMAL-c4x::halA]* was created by transforming BL21(DE3) with the eJM68 plasmid, cloned from a Gibson assembly consisting of 2 fragments: 1) *halA* fragment: PCR with primers oJM229 and oJM230 from DS2 gDNA template; 2) ampR-ori_f1-MBP fragment: PCR with primers oJM227 and oJM228 from pMAL-c4x (20) plasmid DNA template.

*eJM118 [BL21(DE3) pMAL-c4x::bhalB]* was created by transforming BL21(DE3) with the eJM117 plasmid, cloned from a Gibson assembly consisting of 2 fragments: 1) *bhalB* fragment: PCR with primers oJM259 and oJM260 from a synthetic DNA template; 2) ampR-ori_f1 fragment: PCR with primers oJM227 and oJM233 from pMAL-c4x (20) plasmid DNA template.

**Growth Curves**

*Hfx. volcanii* strains were grown as described above at 42°C until an $OD_{600nm}$=0.5 and diluted 10-fold to an $OD_{600nm}$=0.05. 200µl of diluted culture was transferred to a 96-well flat-bottom plate (Corning Inc., #3370) next to Hv-Cab blanks. All wells surrounding the samples were filled with 200µl $ddH_2O$ to prevent media evaporation. Growth curves of biological triplicates were performed using an EPOCH2 microplate spectrophotometer (Agilent) with constant orbital shaking at 42°C. $OD_{600nm}$ measurements were collected every 30 minutes for 48 hours. Each measurement was averaged across triplicates and then subtracted from the averaged Hv-Cab blank measurements. Biological replicates were averaged, the standard error of the mean was calculated, and an exponential equation was plotted using libraries in Python (Pandas v.1.5.3 and Matplotlib v.3.7.1, as previously cited).

**Spot Dilution Viability Assay**

*Hfx. volcanii* strains were grown at 42°C until an $OD_{600nm}$=0.5 and were diluted to an $OD_{600nm}$=0.2, then serially diluted 10-fold. 30 µl of the serial dilutions were spotted onto standard Hv-Cab 2% agar plates. Plates were left covered at room temperature to dry overnight and placed in plastic bags at 45ºC for 2-3 days until single colonies were observed. Pictures of plates were taken after 5-7 days at room temperature.

**Plate Motility Assay**

Hv-Cab 0.3% agar plates were poured (20 ml per plate) fresh on the day of inoculation and cooled for 15 minutes at room temperature. A 1µl droplet of mid-exponential culture was pipetted on top of the agar in each of the four quadrants of the plate, where mutant strains were inoculated with the parental wild-type strain on the same plate. Plates were incubated face-up in a plastic bag at 37°C. All bags were placed within a closed plastic container, and a slightly damp microfiber cloth was placed on top of each bag. Pictures were taken after four days of growth. The area of the motility halos was measured manually using Fiji, and the motility of each mutant was normalized by their respective wild-type measurements from the same plate.

**Western Blots**

*Hfx. volcanii* cells were grown to exponential phase ($O.D._{600}$=0.5). 2mL of culture were pelleted by centrifugation (4000xg, 2 minutes at room temp), supernatant removed and stored at -80°C. Cell pellets were resuspended using 50uL 5X-SDS protein running buffer solution (10% w/v SDS, 5% v/v 2-Mercaptoethanol, 30% v/v glycerol, 250mM Tris pH 6.8, bromophenol blue) and 50uL dH20. Samples were boiled at 95℃ for 10 minutes. 10uL of each sample were loaded into an Invitrogen 4-12% NuPAGE Bis-Tris protein gel and run at 200V for 35 minutes in 1X MES buffer. Samples were transferred to a nitrocellulose membrane (Amersham Protran) via a BioRad Trans-Blot Turbo Transfer System. The membrane was incubated in blocking solution (5% milk in 1X PBST) for 45 minutes at room temperature. Following blocking the membrane was incubated in primary anti-GFP monoclonal mouse antibody (1:$10^3$ dilution) overnight at 4°C on a rocking platform. Primary antibody was removed by three 5-minute washes in 1X PBST followed by an incubation with secondary anti-mouse antibody in 1X PBST (1:$10^4$ dilution) for 45 minutes at room temperature on a rocking platform. Secondary was removed vi a 15-minute wash with 1X PBST and three subsequent 5-minute washes. Membranes were Imaged on BioRad ChemiDoc MP.

**Expression and Purification of HalA and bHalB**

Cells were grown at 37°C in 2xYT media (Sigma-Aldrich) supplemented with carbenicillin to an $O.D._{600}$ of 0.6 and induced with 1 mM IPTG. Temperature was kept at 37°C for 4 hours and then cells were harvested at 1400 xg for 30 minutes at 4°C. Cells were resuspended in lysis buffer (1 M NaCl, 50 mM CHES pH=10) supplemented with 5 mM EDTA and 100 µM PMSF and lysed by sonication using a 550 Sonic Dismembrator (Fisher Scientific). Sonication was done on ice with pulses of 30 seconds at 15% power with 30 seconds of rest for a total pulse duration of 5 minutes. Insoluble material was pelleted at 18,000 xg for 30 minutes at 4°C and soluble lysate was collected. Water was then added slowly with gentle agitation to make a 1:6 dilution (final concentrations of 167 mM NaCl, 8 mM CHES pH=10). For HalA preparations, samples were incubated with 2 ml of amylose resin (New England Biolabs) for 1 hour at 4°C with gentle agitation. The resin was collected and washed in a 50 ml falcon tube with 15 ml of low salt buffer (167 mM

NaCl, 8 mM CHES pH=10) three times. The resin was then incubated with elution buffer (167 mM NaCl, 8 mM CHES pH=10, 10 mM Maltose) for 10 minutes at 4°C. The sample was then spun in a benchtop centrifuge at 3000 rpm for 5 minutes at 4°C to pellet the resin. The supernatant was collected and allowed to incubate overnight with PreScission protease with gentle agitation. Samples were then adjusted to pH 7 by the addition of 1 M Tris pH=7.4 to a final concentration of 20 mM after dropping the water dilution step. Samples were stored at 4°C until ready for electron microscopy preparation. For bHalB preparations, lysates were neutralized by adding 1M Tris pH=7.4 to a final concentration of 20mM and stored at 4°C until ready for electron microscopy preparation.

**Negative Staining Electron Microscopy**

4 μl of the purified protein samples was applied to glow discharged copper mesh Formvar coated carbon grids and were allowed to adsorb to the grid for about 30 seconds. The grids were briefly washed in double distilled water, followed by staining with 2.5% Aqueous Uranyl Acetate, and allowed to dry fully before imaging. Grids were imaged in a FEI Morgagni transmission electron microscope (FEI, Hillsboro, OR) operating at 80 kV and equipped with a Nanosprint5 CMOS camera (AMT, Woburn, MA).

**Live-Cell Imaging and Analysis**

*Snapshots:* Phase-contrast and fluorescence microscopy snapshots were taken from cultures concentrated 10-fold by centrifuging (3,000xg for 2 minutes), and 3μl droplets of culture were placed on 60×24 mm coverslips and gently immobilized under pre-warmed 1.5x0.5 cm, round 0.5% Hv-Cab agarose pad (SeaKem LE Agarose, Lonza, #50002). Phase-contrast snapshots were automatedly segmented using CellPose2 (https://www.cellpose.org/) (21). For accuracy across different cell morphologies and shape defects, a custom trained CellPose2 model (21) was used based on existing datasets in the Bisson Lab unrelated to the data in this work. Masks and ROI files were exported from CellPose2, and cell area, solidity, length, and width were calculated in Fiji (22). Rods were automatically assigned for cells above the arbitrary aspect ratio cutoff of 2.0, while disks were assigned to cells 1<AR<2. Distributions were graphed using PlotsOfData (23).

*HaloTag labeling*: Cells expressing HalA- and HalB-HaloTag fusions were incubated with HaloTag-ligand JF549 and Alexa488 for 30 minutes under optimal growth conditions (agitation at 42°C). Cells were harvested, and samples were prepared as described above. However, in this case, coverslips were cleaned by our protocol for single-molecule imaging (19). Briefly, coverslips were placed in "zigzag" orientation in a custom 3D-printed tray, sonicated for one hour in 2% Hellmanex III detergent (Hellma, #9-307-011-4-507), rinsed twice with Milli-Q water, sonicated another hour in 190 proof, benzene-free ethanol, rinsed twice with ethanol and parafilm-wrapped stored in fresh ethanol.

*HalA-msGFP and HalB-msfGFP Time-Lapses:* The same protocol as above was followed for time-lapses shown in Fig. 2, but cells were immobilized under a pre-warmed 2.5x0.5 cm round 0.3% agarose pad in a 50 mm Mattek dish (MatTek Corp., #P50G-1.5-30-F). Images were collected every 2 minutes for 24 hours. Polymerization and depolymerization events were randomly chosen, and kymographs were created manually using the KymographBuilder plugin (https://github.com/fiji/KymographBuilder) in Fiji. Rates were calculated from the "simple linear regression" slope in time versus filament length curves in GraphPad Prism 10 (Dotmatics, https://www.graphpad.com).

*Solidity Tracking during Cell Division, Growth, and Shape Transition:* Fresh colonies were washed out and resuspended in 500 μL of media directly from Hv-Cab plates, then pelleted (3,000xg for 2 minutes), resuspended in 100uL of media and placed onto a clean 50 mm Mattek dish. Cells were immobilized under a pre-warmed 2.5x0.5 cm round 2.5% Hv-Cab agarose pad. Images were collected from several XY positions every 2 minutes for 12-24 hours. Images were then cropped, selecting arbitrary events of cytokinesis (start of the division cleft until daughter cells' separation), elongation (from cell birth to onset of cytokinesis), and shape shifts (disk-to-rod transitions). To avoid issues with defect propagation, only cells with initial area and solidity values within the coefficient of variance intervals measured from the wild-type population were chosen for analysis. Crops were combined as stacks, segmented by Omnipose (Cutler) trained with our group's phase-contrast microscopy image collection from different strains. Binary masks were then imported to the plugin MicrobeJ (24) in Fiji, extracting solidity over time. Data were plotted using PlotTwist (25).

All live-cell imaging described above was imaged at 42°C using a Nikon TI-2 Inverted Microscope within an Okolab H201 enclosure. Phase-contrast and GFP-fluorescence images were acquired with a Hamamatsu ORCA Flash 4.0 v3 sCMOS Camera (6.5 µm/pixel), a CFI PlanApo Lambda 100x DM Ph3 Objective, and a Lumencor Sola II Fluorescent LED (380–760 nm).

*3D SoRa Super-Resolution:* Cells were stained with 300µM Nile Blue A (Sigma, #A17174), and a 3µl droplet of culture was immediately transferred to 35mm glass bottom dishes (Ibidi, #1218-200) and immobilized under a 1.5x0.5 cm, round 0.3% Hv-Cab agarose pad. Cells were imaged at room temperature using a Nikon Ti-2 (Nikon) equipped with a Yokogawa CSU-W1 SoRa spinning disk, with 561 nm (Nile Blue A) and 488 nm (GFP) lasers, equipped with a Plan Apo λ 100x objective, Prime BSI-Express camera (pixel size: 0.02 µm), with 41 Z-slices of 0.1µm. 3D stacks were processed using NIS-Elements software (Nikon), using NIS.ai for denoising and 3D deconvolution. Stacks were then rendered into 3D projections using Fiji.

*Cell Surface Curvatures:* For the cumulative distributions of membrane curvature, 176 cells were analyzed (77 rods and 99 disks) from the wild-type strain, 191 cells (97 rods and 94 disks) from the Δ*halA* strain, 217 cells (83 rods and 134 disks) from the Δ*halB* strain, and 174 cells (91 rods and 83 disks) from the Δ*halAB* strain. Crops of individual membrane-labeled cells of 3D SoRa images were combined as stacks and fed into Fiji's LimeSeg plugin (26). Settings were set as D_0 to 4, F_pressure to 0.025, Z_scale to 4.307, Range_in_d0_units to 5, and RealXYPixelSize to 0.0232. Vertices of the triangular mesh were extracted, and the associated Gaussian curvature was calculated at those points. Gaussian curvature was converted from pixel units to um units using the xy pixel size 0.0232um/pixel. 3D colormap projections were generated using a custom Python script (https://github.com/Archaea-Lab/halofilinScripts). The ".ply" files from LimeSeg containing the vertex, edge, and face information were imported into Blender (https://www.blender.org/). The Gaussian curvature colormap was then mapped onto the cell using another custom Python script (https://github.com/Archaea-Lab/halofilinScripts) within the blender text editor. The colormap, now stored as a "color attribute" of the cell, was then placed on a new material so that the cell could be placed in a camera view and an image rendered. Cumulative distributions were calculated, curves fit-spline (20-point bins) and graphed using GraphPad Prism 10.

*Mapping Cell Surface Curvature to Halofilin Location:* For the correlation between HalA- and HalB-msfGFP fluorescence signal and membrane curvature, 49 cells were analyzed (26 rods and 23 disks) of HalA-msfGFP (strain aJK3) and 35 cells (22 rods and 13 disks) of HalB-msfGFP (strain aBL404). A custom Python script (https://github.com/Archaea-Lab/halofilinScripts) was used to analyze the location of halofilin molecules based on Gaussian curvature data. First, the background subtracted pixels' grey values from the 3D-SoRa's cell membrane images and registered them with their XYZ location. Next, the grey values of the XYZ positions were matched with the Gaussian curvature from the LimeSeg analysis. Grey values were then normalized by the frequency amount of each curvature grouped in bins of 10. Data were plotted using PlotTwist.

*Halofilins and Z-ring Positions across rod cells:* Phase-contrast Images of cells expressing HalA-msfGFP, HalB-msfGFP, FtsZ1-msfGFP or FtsZ2-msfGFP were segmented by CellPose2 as described above. A medial axis of rod-shaped cells was then traced from binary masks using Fiji's MicrobeJ plugin. From the medial axis, intensity profiles were traced, and the median point of each Gaussian fitted curve was used as the Z-ring position. This dataset was graphed against each respective cell's area, generating the plot in Figure 5C. Moreover, using the same strategy described for Gaussian curvatures, the HalB-msfGFP, FtsZ1-msfGFP and FtsZ2-msfGFP grey values in each pixel were correlated to the position of the medial axis of each rod cell to generate the heatmaps in Figure 5B. Data were plotted using PlotTwist.

*Hfx. volcanii Spheroplasting:* Cells were spheroplasted as described in Cline et al, 1989 (27). Briefly, 2 mL of cell culture was centrifuged at 4500xg for 10 minutes, resuspended in 2 mL Buffered Spheroplasting Solution, centrifuged again at 4500xg for 10 minutes, and resuspended in 200uL of Buffered Spheroplasting Solution. A 20uL drop of 0.5M EDTA was added, and cells were incubated at room temperature for 5 minutes. For SoRa imaging, cells were stained with 300 µM Brilliant Blue (Sigma). Images were collected as described above.

*HalA-msfGFP filament length and HalB-msfGFP foci-counting measurements:* Phase-contrast and fluorescence snapshots were automatedly segmented using a custom trained Omnipose model (28). The model was trained based on previously existing datasets in the Bisson Lab. Masks and ROI files were exported from Omnipose, and foci were linked to their parent cells using CellProfiler (29). The analysis pipeline can be found on the Bisson Lab GitHub: https://github.com/Archaea-Lab/Omnipose_Cellprofiler-Pipeline).

**Supplemental Figures**

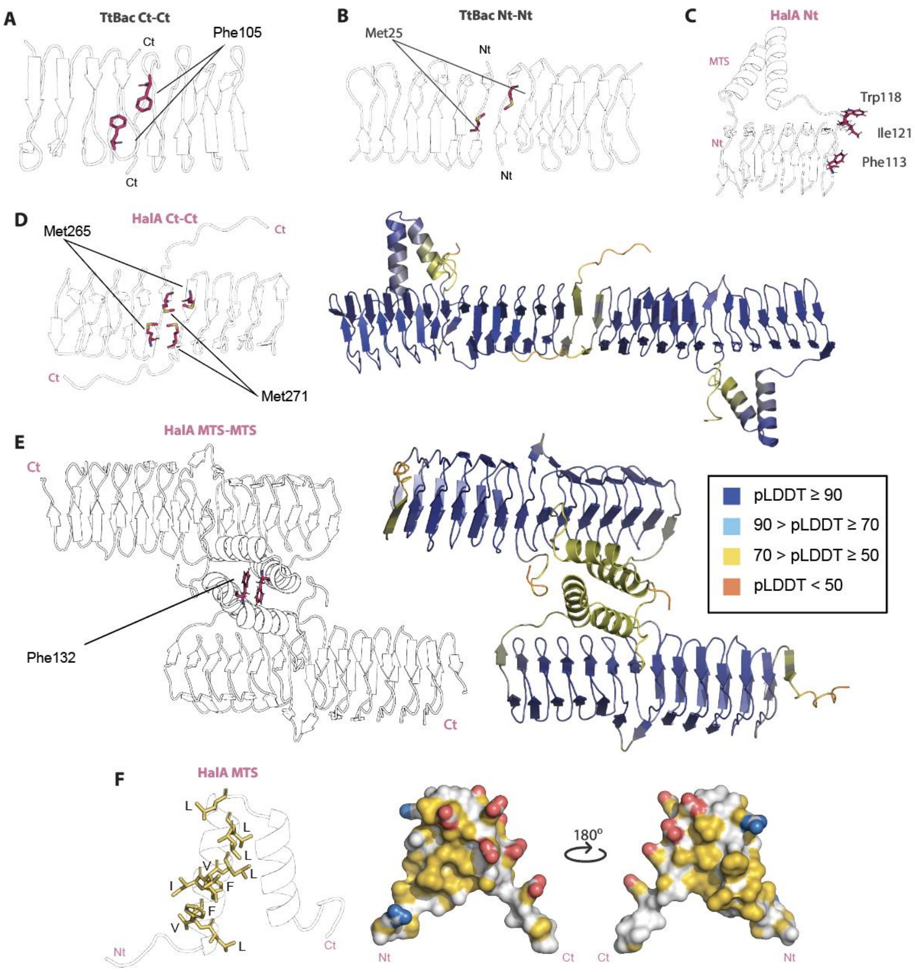


**Figure S1:** AlphaFold2 models of Haloferax volcanii HalA predicting filamentation interfaces. Cryo-EM structure of TtBac (PDB 6rib) highlighting the (A) C-terminus:C-terminus and (B) N-terminus:N-terminus polymerization interfaces. Phenylalanine 105 and Methionine 25 were shown to be relevant for the Ct:Ct and Nt:Nt interfaces, respectively. (C) AlphaFold2 model of Hfx. volcanii HalA highlighting its N-terminus surface of the bactofilin. AlphaFold2 did not predict a dimerization interface. F113, W118, and I121 residues are oriented into solvent space. (D) AlphaFold2 model of a homodimer of HalA (pTM = 0.61, ipTM = 0.34) highlighting the Methionines 265 and 271 of HalA, which appear to come in close contact with each other, comparable to TtBac's F105. (E) AlphaFold2 model of a homodimer of HalA highlighting a possible MTS:MTS polymerization interface (pTM = 0.49, ipTM = 0.1299). Phenylalanines 132 form an aromatic stacking interaction matching the TtBac interfaces. (F) AlphaFold2 model highlighting the hydrophobic residues on the surface of the first helix of the HalA MTS hairpin (left) and the surface hydrophobicity (right). Yellow: polar residues; Red: negatively charged residues; Blue: positively charged residues; White: all remaining atoms, including the polar backbone. AlphaFold2 models are colored according to the per-residue model confidence score (pLDDT). Hydrophobicity surface maps were created using the YRB Pymol script (30).

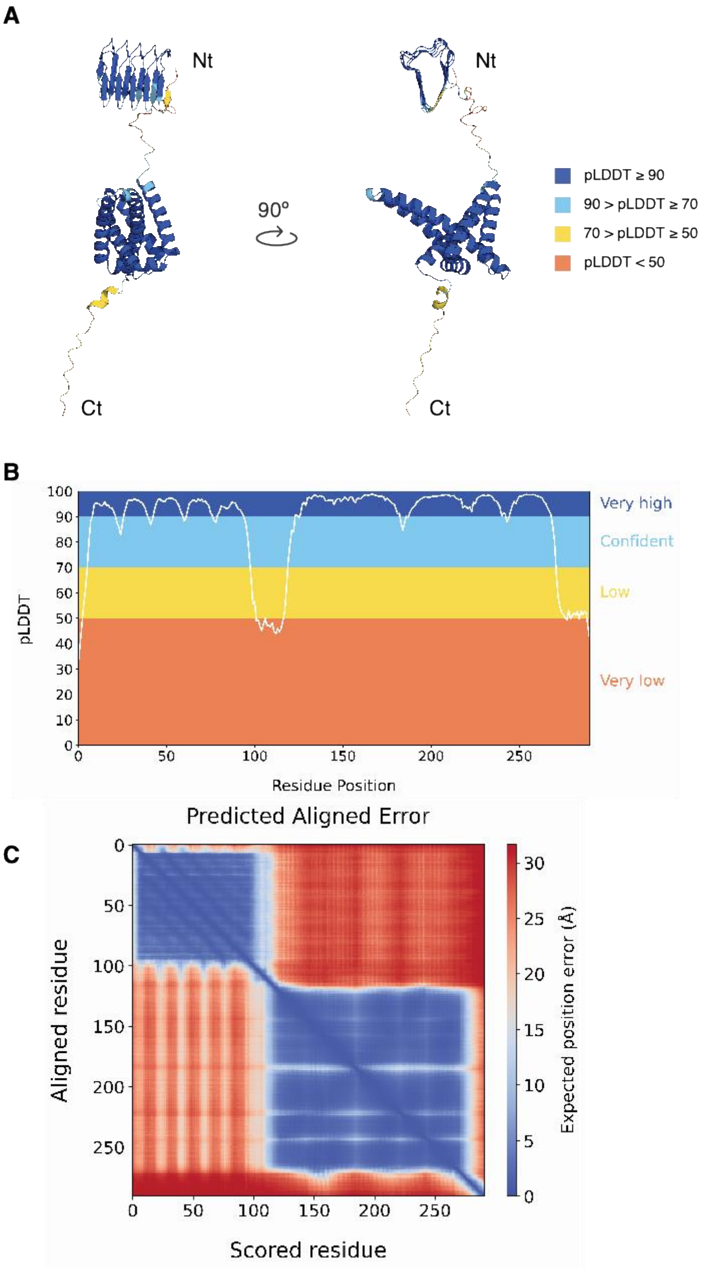


**Figure S2: (A)** Predicted structure of HalB from *Hfx. volcanii*, colored according to the per-residue model confidence score (pLDDT) produced by AlphaFold2. Structures lack the signal peptide (see Methods). **(B)** Distribution of pLDDT values in function of the residue position of the protein sequence of HalB. The plot follows the same color scheme as in sub-panel (A). **(C)** Predicted Aligned Error (PAE) plot for the predicted structure of HalB.

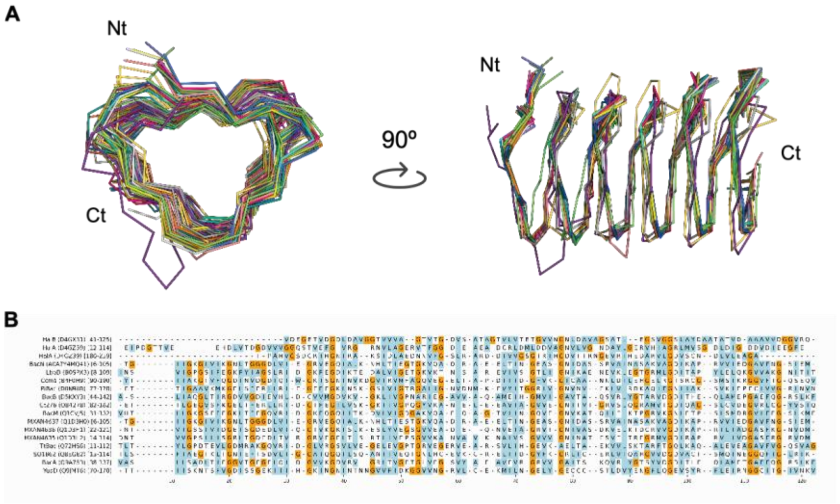


**Figure S3:** Multiple structure alignment **(A)**, and structure-guided multiple sequence alignment **(B)** of halofilin and bactofilin domains. Both structure and sequence alignments contain the N- and C-terminal bactofilin domains of HalA, bHalB, and the bactofilin domains of 14 canonical bactofilins (see Table S1 for more information). Each sequence is preceded by its protein name, the UniProt ID, and the residue coordinates of its bactofilin domain, respectively. Glycines are highlighted in orange, and hydrophobic residues are highlighted in blue.

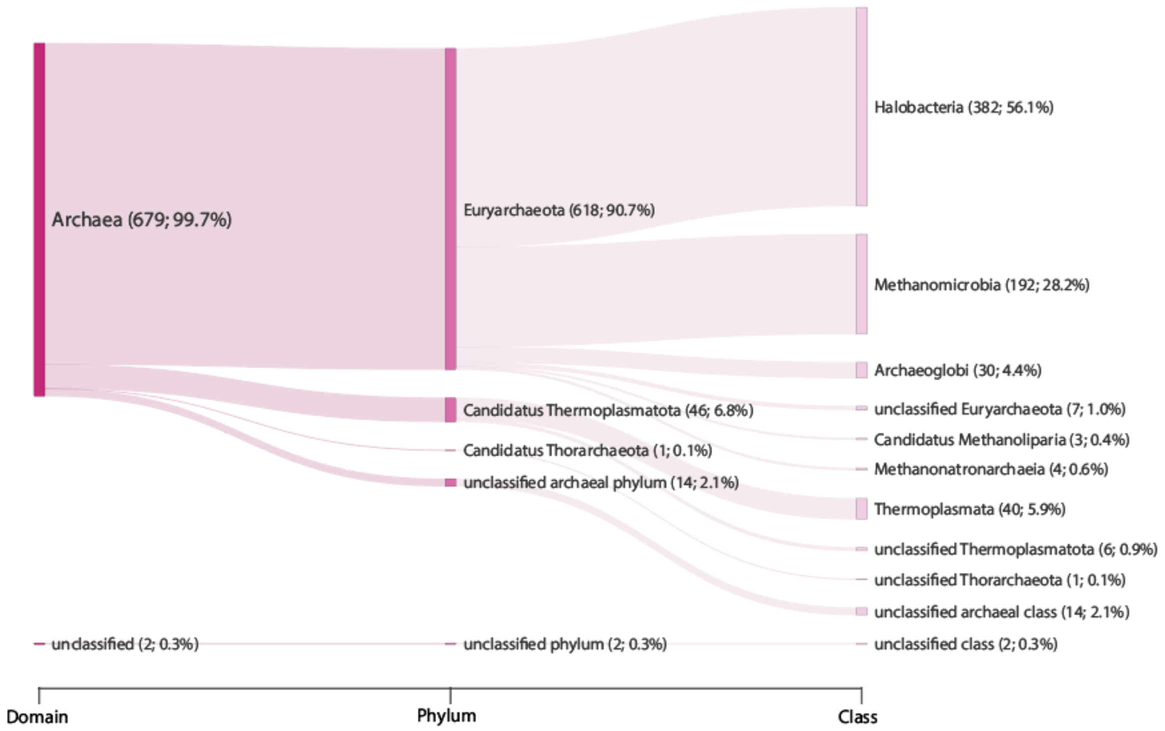


**Figure S4:** Taxonomic representation of the protein sequences issued from the HalA cluster from Figure 1A. Each node is annotated with the number of sequences issued from that taxonomic rank and corresponding percentage. Every percentage was calculated concerning the total number of sequences in the cluster. Ranks missing from the NCBI Taxonomy database are either labeled as or prefixed by "unclassified".

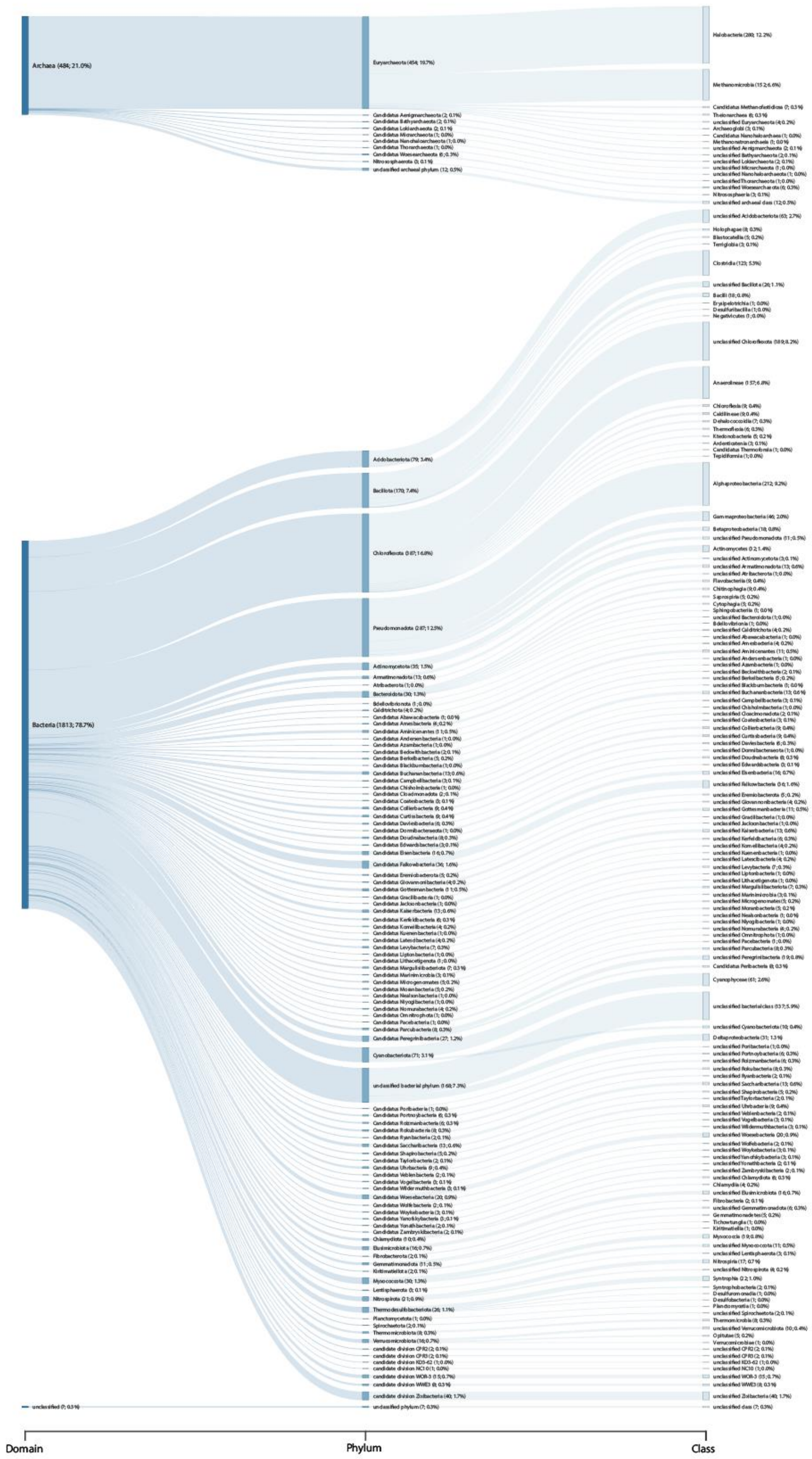


**Figure S5:** Taxonomic representation of the protein sequences issuing from the HalB cluster from Figure 1A. Each node is annotated with the number of sequences issued from that taxonomic rank and corresponding percentage. Every percentage was calculated concerning the total number of sequences in the cluster. Ranks missing from the NCBI Taxonomy database are either labeled as or prefixed by "unclassified".

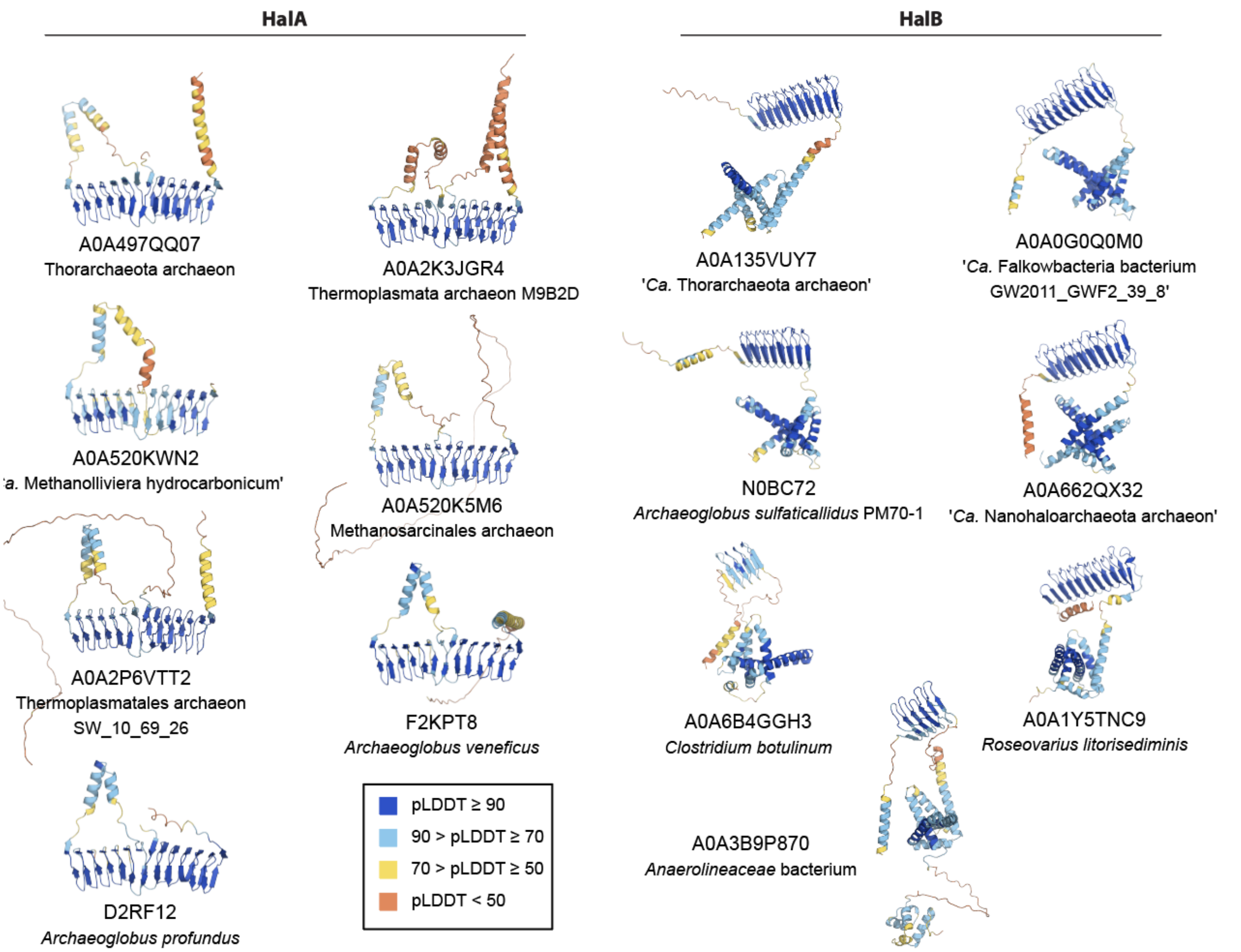


**Figure S6:** AlphaFold2 models for HalA and HalB structural variants. These models correspond to the sequences used to seed the HalA and HalB clusters from Figure 1A (see Table S1). All models were downloaded from the AlphaFold Protein Structure Database (31), and each is annotated with its respective accession. The accessions are shared by both UniProtKB and the AlphaFold Protein Structure Database. Each model is colored according to the per-residue model confidence score (pLDDT) produced by AlphaFold2.

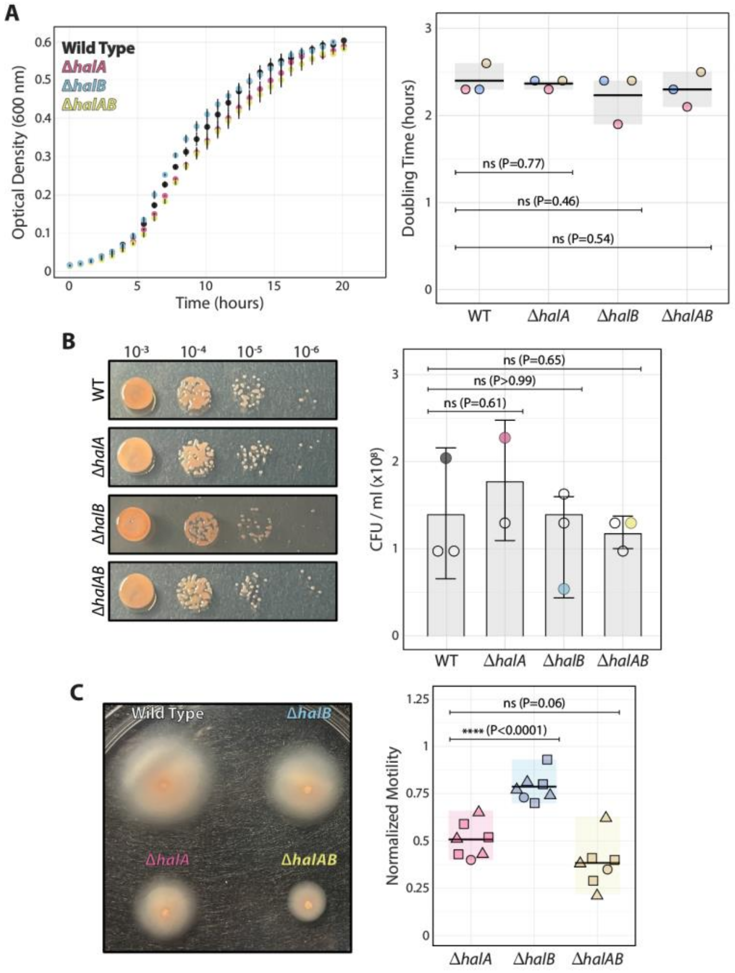


**Figure S7:** Halofilin mutants show motility but not viability defects. **(A)** Bulk growth curves of wild type, Δ*halA*, Δ*halB,* and Δ*halAB* strains. Mutant doubling times (2.3±0.1, 2.2±0.3, and 2.3±0.2, respectively; mean±SD) were not statistically different (P-values from unpaired t-student tests shown in graph) compared to wild type (2.4±0.2; mean±SD). **(B)** Spot dilution viability assay of wild type, Δ*halA*, Δ*halB,* and Δ*halAB* strains. Colony formation unit (CFU/ml x $10^{-8}$) from mutants (1.22±0.71, 1.11±0.69, and 1.22±0.19, respectively; mean±SD) were not statistically different (P-values from unpaired t-student tests shown in graph) compared to wild type (1.44±0.77; mean±SD). Colored datapoints in the bar plot (right) correspond to the spot dilution images (left). **(C)** Low-agar (0.3%) motility plates as a proxy for swimming behavior. Motility quantification was done by measuring the halo area from segmented images in Fiji. Each datapoint in the boxplot represents technical replicates (same geometrical shape) over biological replicates (different geometrical shapes). Motility of Δ*halA* mutant (0.51±0.1; mean±SD) showed significant differences (P-values from unpaired t-student tests shown in graph) from Δ*halB* (0.79±0.1; mean±SD), but not Δ*halAB* (0.38±0.1; mean±SD).

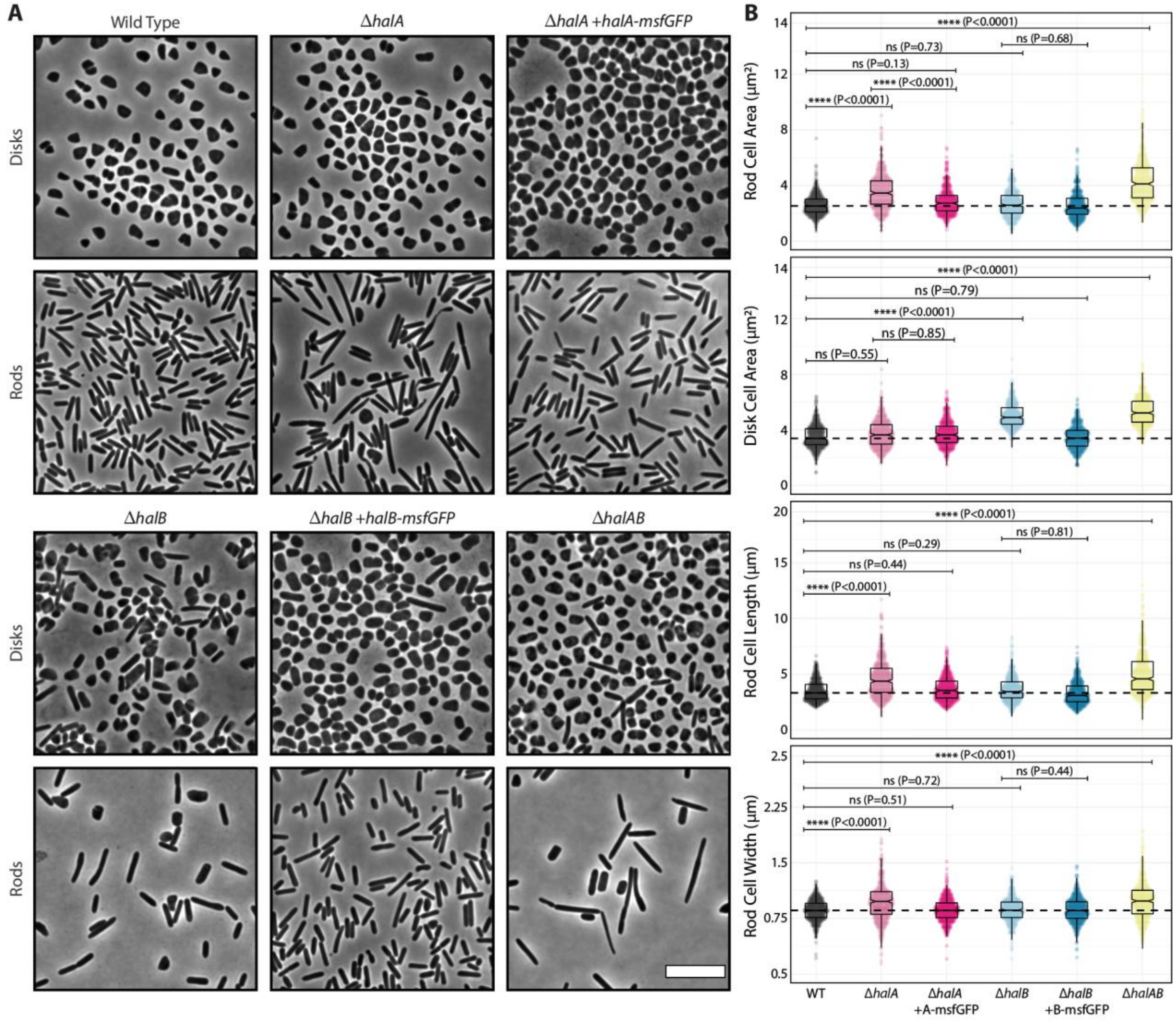


**Figure S8:** Quantitative description of cell morphology and size from halofilin mutants. **(A)** Phase contrast microscopy of the deletion strains shows morphological defects in rod- and disk-cell types. **(B)** Quantification of each strain's cell area, rod cell length, and rod-cell width. Kolmogorov-Smirnov tests were applied to non-parametric pairs between wild-type and every other sample, resulting in statistically differences labeled with their associated P-values, and non-significant comparison noted with "ns". Dashed lines indicate the mean of the wild type distributions. Scale bars represent 5μm.

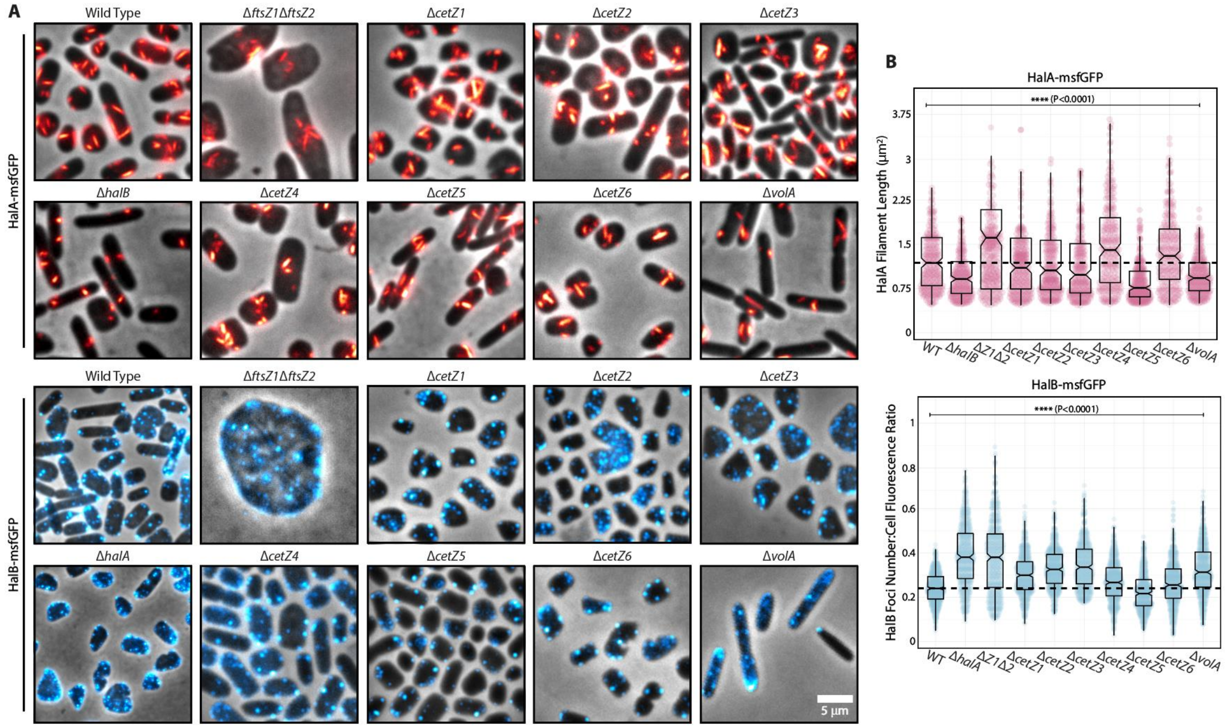


**Figure S9:** HalA-msfGFP and HalB assembly is independent of other cytoskeletal systems **(A)** HalA-msfGFP and HalB-msfGFP live-cell imaging across a variety of cytoskeleton mutants backgrounds. **(B)** HalA length and HalB foci number quantitation. Kolmogorov-Smirnov tests were applied to non-parametric pairs between wild-type and every other sample, resulting in significant differences (P<0.0001).

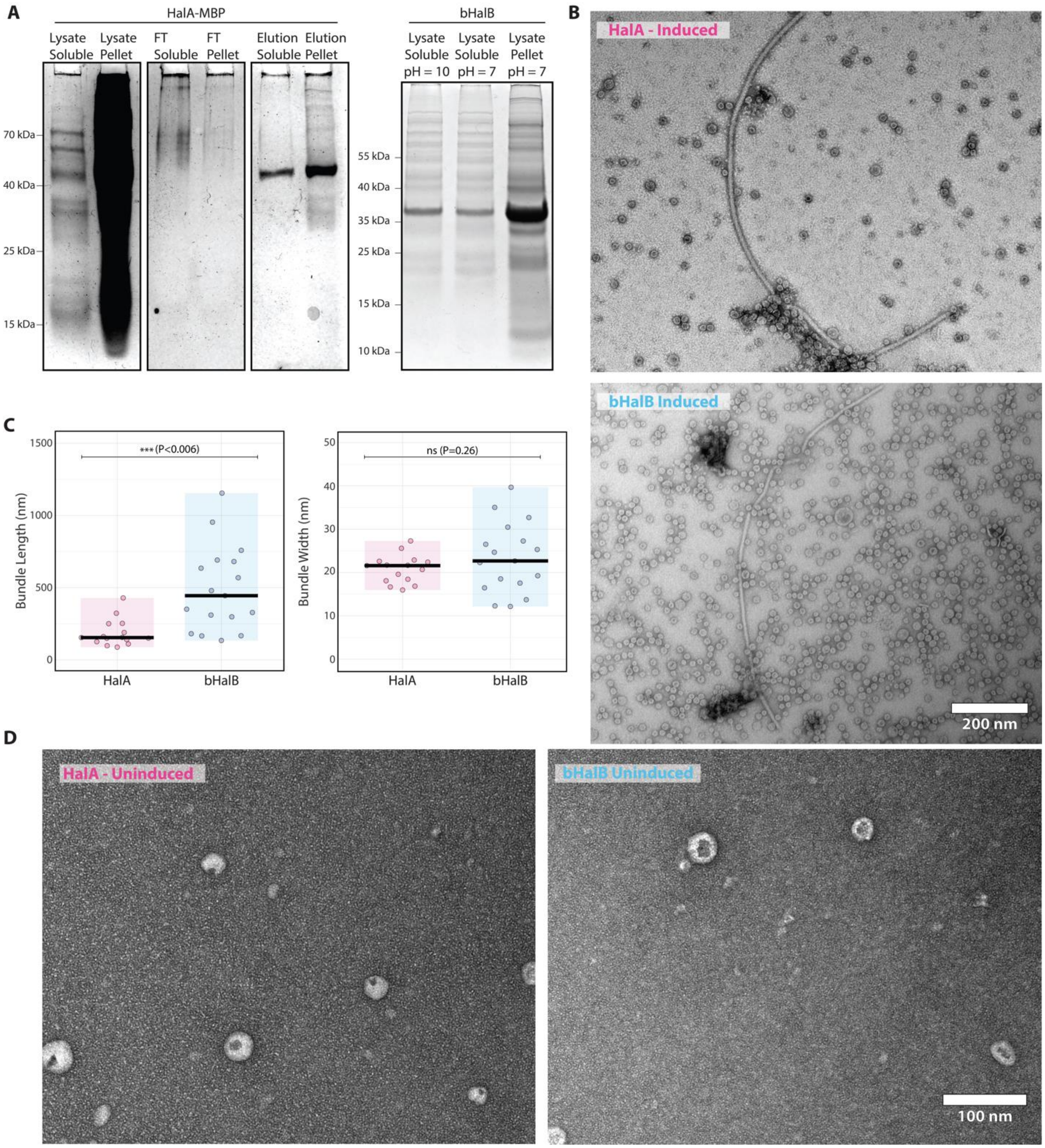


**Figure S10:** Purification and *in vitro* imaging of HalA and bHalB **(A)** purification SDS-PAGE gels of each purification step of the HalA (left) and bHalB (right) samples detailed in methods. **(B)** TEM images of HalA and bHalB bundles at low magnification. **(C)** HalA and bHalB bundles, despite having similar appearance, they show significant differences in length, but not width. ANOVA tests were applied, and P-values shown in graphs and non-significant comparison noted with "ns". **(D)** TEM images from uninduced HalA and bHalB fractions. Similar toroid structures were observed as in the purified samples, but more scarce and smaller compared to those in the induced fraction.

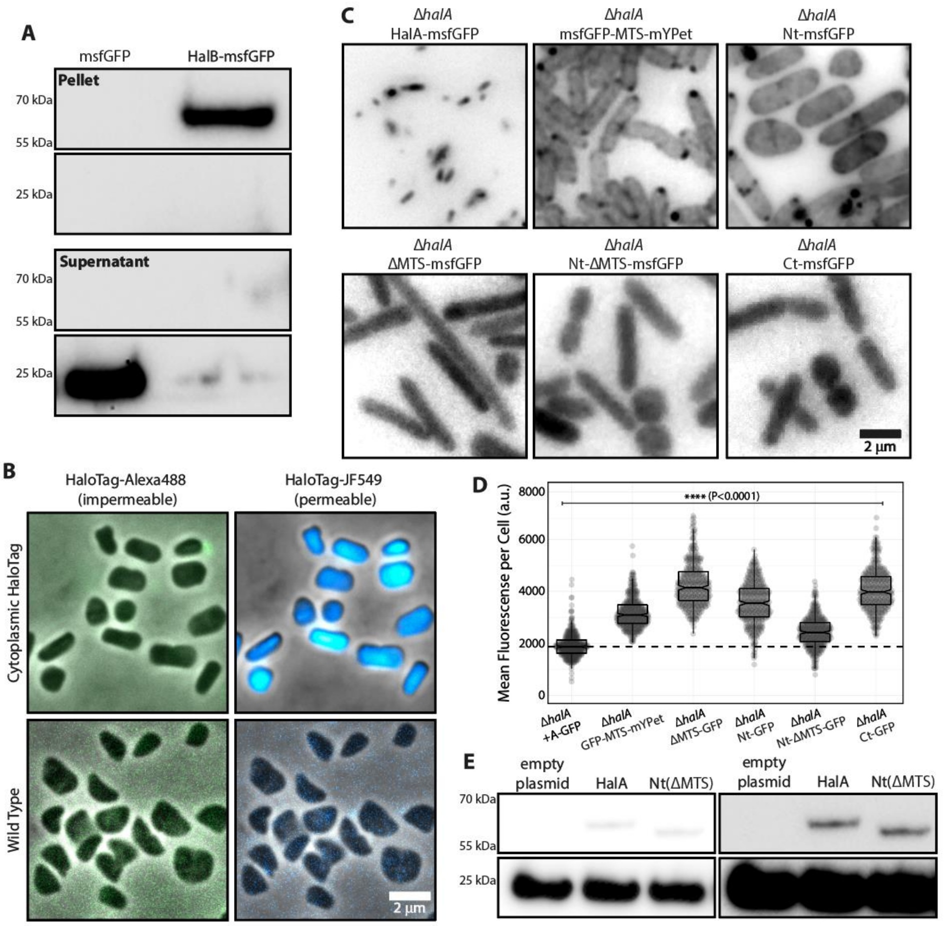


**Figure S11: (A)** Western blot of cell lysates expressing cytoplasmic msfGFP (left) and HalB-msfGFP (right) from insoluble (top) and soluble (bottom) fractions. Cells were lysate as described in methods and fractionated by ultra-centrifugation at 50,000 xg for 30 minutes. **(B)** HaloTag-ligand JF549 is permeable, but not Alexa488, and reacts with cytoplasmic HaloTag. (Bottom) Wild-type cells were simultaneously incubated with HaloTag-ligand JF549 and Alexa488 (1μM each) and imaged by phase contrast and epifluorescence microscopy. (Top) Cells expressing free cytoplasmic HaloTag were again incubated with HaloTag-ligand JF549 and Alexa488 (1μM each) and imaged by phase contrast and epifluorescence microscopy. **(C)** Cells expressing different HalA-msfGFP constructs were expressed under 500 μM tryptophan and imaged by epifluorescence microscopy. **(D)** Quantification of mean grey values from cells expressing different HalA-msfGFP constructs. All samples showed means above and statistically different than the HalA-msfGFP (Kolmogorov-Smirnov non-parametric tests). **(E)** Western blot from lysates of cells constitutively expressing cytoplasmic msfGFP paired with a negative control (left), full-length HalA-msfGFP (center) and the HalA(Nt)(ΔMTS)-msfGFP truncate (right). The left and right panel represent the same image under different contrast levels.

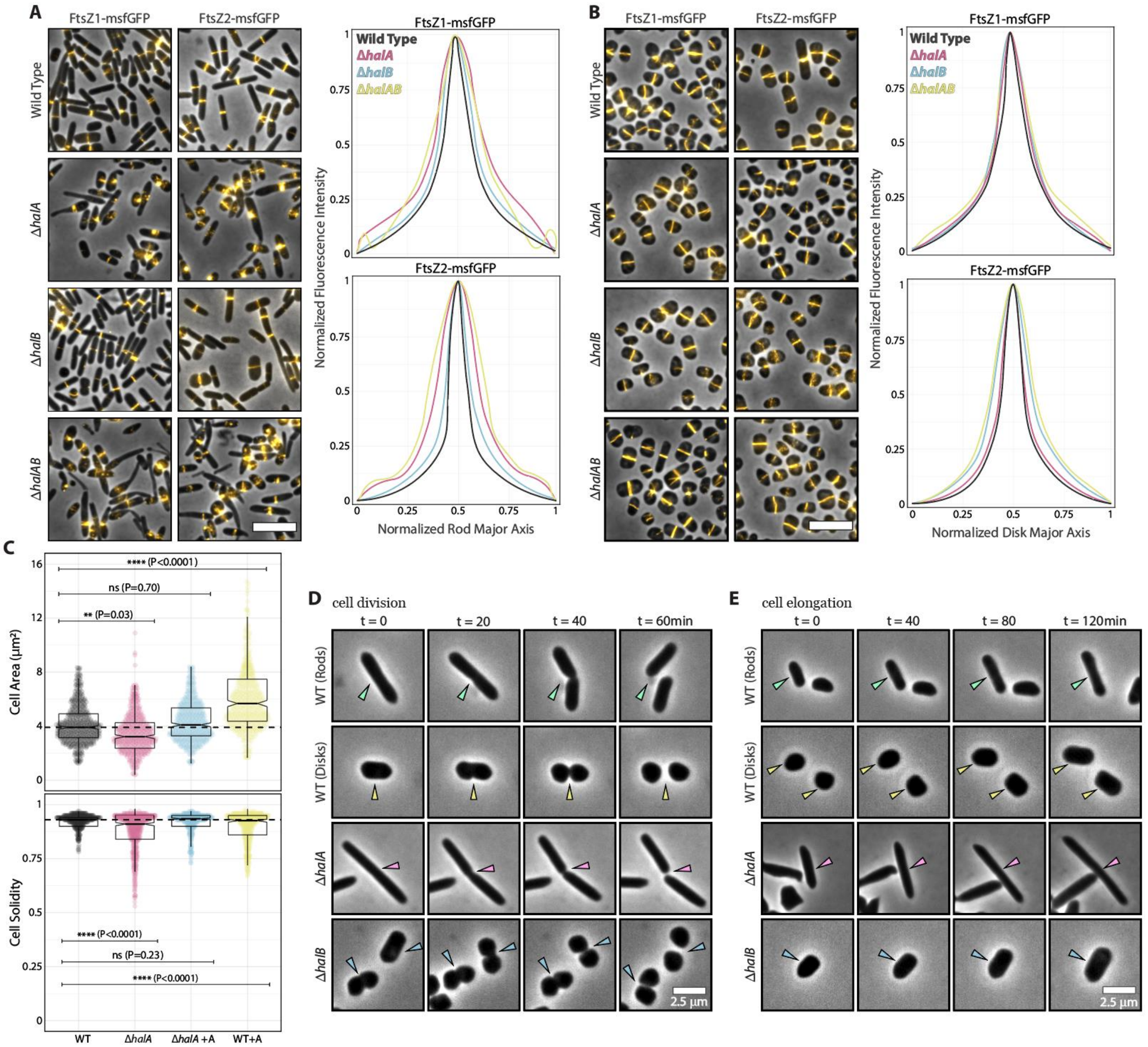


**Figure S12:** Z-ring decondensation in rod mutants does not imply a role of halofilins in cell division and elongation. Phase contrast and epifluorescence overlays show the cellular localization of the cytokinetic Z-ring by labeling the tubulin homologs FtsZ1-msfGFP and FtsZ2-msfGFP in **(A)** rods and **(B)** disks. Normalized fluorescence intensities were plotted against the normalized major axis of the cells. **(C)** *Hbt. salinarum* cell area and solidity were measured from phase contrast images represented in Fig. 5C. Kolmogorov-Smirnov tests were applied to non-parametric pairs between wild-type and every other sample, resulting in statistically differences labeled with their associated P-value, and non-significant comparison noted with "ns". Representative montages of phase contrast time-lapses with cells undergoing cell division **(D)** and elongation **(E)** within a cell cycle period. Unless specified, scale bars represent 5μm.

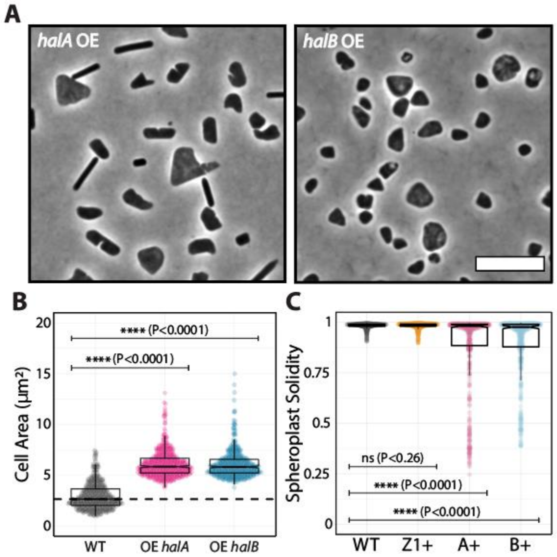


**Figure S13:** High expression levels of *halA* and *halB* result in morphological defects. **(A)** Phase contrast microscopy of cells overexpressing *halA* and *halB*. **(B)** Quantification of cell area from overexpression strains. **(C)** Quantification of solidity of spheroplasted cells overexpressing *ftsZ1*, *halA* and *halB*. Kolmogorov-Smirnov tests were applied to non-parametric pairs between wild-type and every other sample, resulting in statistically differences labeled with their associated P-value, and non-significant comparison noted with "ns". Scale bars represent 5µm.

**TABLE S1:** Metadata for the protein sequences used as seeds in the cluster map's BLASTp searches

| Canonical bactofilin seeds | | | | |
|---|---|---|---|---|
| **ID** | **Name** | **Organism** | **Database** | **Reference** |
| Q9A753 | BacA | *Caulobacter crescentus* | UniProt | (32) |
| D5KXY3 | BacB | *Caulobacter crescentus* | UniProt | (33) |
| Q1CVJ5 | BacM | *Myxococcus xanthus* | UniProt | (32) |
| A0A7Y4MQ41 | BacN | *Myxococcus xanthus* | UniProt | (34) |
| A0A8E4SGC6 | BacO | *Myxococcus xanthus* | UniProt | (34) |
| A0A7Y4JG82 | BacP | *Myxococcus xanthus* | UniProt | (34) |
| B4F0H9 | Ccm1 | *Proteus mirabilis* | UniProt | (32) |
| B0SPX3 | LbbD | *Leptospira biflexa* | UniProt | (35) |
| ACI28222.1 | HPG27_1480 | *Helicobacter pylori* | GenBank | (32) |
| Q72HS6 | TtBac | *Thermus thermophilus* | UniProt | (32) |
| D0N980 | PiBac | *Phytophthora infestans* | UniProt | (32) |
| AAD08589.1 | HP1542 | *Helicobacter pylori* | GenBank | (36) |
| Q8EGE2 | SO1662 | *Shewanella oneidensis* | UniProt | (33) |
| Q1D3H2 | MXAN4635 | *Myxococcus xanthus* | UniProt | (33) |
| Q1D3H1 | MXAN4636 | *Myxococcus xanthus* | UniProt | (33) |
| Q1D3H0 | MXAN4637 | *Myxococcus xanthus* | UniProt | (33) |
| RCW25186.1 | CcmA | *Vibrio parahaemolyticus* | GenBank | (32) |
| P39132 | BacE | *Bacillus subtilis* | UniProt | (37) |

| | | | | |
|---|---|---|---|---|
| Q9JMT6 | YuaD | *Escherichia coli* | UniProt | - |
| O84278 | Ct276 | *Chlamydia trachomatis* | UniProt | (38) |
| **Halofilin A seeds** | | | | |
| **ID** | **Name** | **Organism** | **Database** | **Reference** |
| D4GZ39 | HalA | *Haloferax volcanii* | UniProt | - |
| A0A497QQ07 | na | Thorarchaeota archaeon | UniProt | - |
| A0A2K3JGR4 | na | Thermoplasmata archaeon M9B2D | UniProt | - |
| A0A520KWN2 | na | '*Candidatus* Methanolliviera hydrocarbonicum' | UniProt | - |
| A0A520K5M6 | na | Methanosarcinales archaeon | UniProt | - |
| A0A2P6VTT2 | na | Thermoplasmatales archaeon SW_10_69_26 | UniProt | - |
| F2KPT8 | na | *Archaeoglobus veneficus* | UniProt | - |
| D2RF12 | na | *Archaeoglobus profundus* | UniProt | - |
| **Halofilin B seeds** | | | | |
| **ID** | **Name** | **Organism** | **Database** | **Reference** |
| D4GX31 | HalB | *Haloferax volcanii* | UniProt | - |
| A0A135VUY7 | na | '*Candidatus* Thorarchaeota archaeon' | UniProt | - |
| A0A0G0Q0M0 | na | '*Candidatus* Falkowbacteria bacterium GW2011_GWF2_39_8' | UniProt | - |
| N0BC72 | na | *Archaeoglobus sulfaticallidus* PM70-1 | UniProt | - |
| A0A662QX32 | na | '*Candidatus* Nanohaloarchaeota archaeon' | UniProt | - |
| A0A6B4GGH3 | na | *Clostridium botulinum* | UniProt | - |
| A0A1Y5TNC9 | na | *Roseovarius litorisediminis* | UniProt | - |
| A0A3B9P870 | na | Anaerolineaceae bacterium | UniProt | - |

**TABLE S2:** Metadata used to create the structure models

| ID | Name | Organism | Database | Reference |
|---|---|---|---|---|
| 6RIA-A | TtBac | *Thermus thermophilus* | PDB | (32) |
| D0N980 | PiBac | *Phytophthora infestans* | AlphaFold DB | - |
| P39132 | BacE | *Bacillus subtilis* | AlphaFold DB | - |
| D4GZ39 | HalA | *Haloferax volcanii* | AlphaFold DB | - |
| D4GX31 | HalB | *Haloferax volcanii* | AlphaFold DB | - |

**TABLE S3:** Plasmids used in this work

| Alias | Plasmid (Promoter) | Reference |
|---|---|---|
| pTA962 | pTA962 (PtnA) | (39) |
| pTA131 | pTA131 ( - ) | (40) |
| pAL750 | pAL750 (Pxyl) | (16) |
| pHVID95 | pTA962::ftsZ2-GFP | (41) |
| pIDJL40 | pTA962::ftsZ1-GFP | (41) |
| eBL94 | pTA962::halA-40aa-msfGFP | This work |
| eBL227 | pTA962::Pro27-msfGFP | (16) |
| eBL232 | pTA962:: halA(ΔMTS)-40aa-msfGFP | This work |
| eBL271 | pTA962::halB-30aa-HaloTag(Ct) | This work |
| eBL272 | pTA962::halB-HaloTag(SW) | This work |
| eBL299 | pTA962::halA(Nt)-40aa-msfGFP | This work |
| eBL301 | pTA962::halB-40aa-msfGFP | This work |
| eBL307 | pAL750::halA | This work |
| eBL309 | pAL750::halB | This work |
| eBL311 | pTA962::msfGFP-halA(MTS)-mYPet | This work |
| eBL313 | pTA962::halA(Nt)-40aa-msfGFP | This work |
| eBL314 | pTA962::halA(Nt)(ΔMTS)-40aa-msfGFP | This work |
| eBL319 | pTA962::halA-30aa-HaloTag | This work |
| eBL330 | pNBKO7::hsal_halA | This work |
| eKA3 | pTA962::HaloTag | This work |
| eZC69 | pBlueScript II carrying fragments of ~1500 nucleotides of upstream and downstream of *halB* open reading frame | This work |
| eJM53 | pBlue::pyrE2:: P27GFP-mevR | This work |
| eJM68 | pMAL-c4x::MPB-halA | This work |
| eJM117 | pMAL-c4x::bhalB | This work |

**TABLE S4:** Strains used in this work

| Alias | Genotype | Reference |
|---|---|---|
| H26 | Δ*pyrE2* | (40) |
| H53 | Δ*pyrE2* Δ*trpA* | (40) |
| aJK3 | Δ*pyrE2* pTA962::halA-40aa-msfGFP | This work |
| aBL80 | Δ*trpA* Δ*pyrE2* Δ*halA* | This work |
| aBL327 | Δ*trpA* Δ*pyrE2* Δ*halA* pTA962::halA-msfGFP | This work |
| aBL336 | Δ*pyrE2* pTA962::halA-30aa-HaloTag pTA962::halA(ΔMTS) | This work |
| aBL369 | Δ*pyrE2* pTA962::halB-30aa-HaloTag(Ct) | This work |
| aBL370 | Δ*pyrE2* pTA962::halB-30aa-HaloTag(SW) | This work |
| aBL401 | Δ*pyrE2* pTA962::halA-30aa-HaloTag pTA962::halA(Ct) | This work |
| aBL404 | Δ*pyrE2* pTA962::halB-40aa-msfGFP | This work |
| aBL412 | Δ*pyrE2* pAL750::halA | This work |
| aBL414 | Δ*pyrE2* pAL750::halB | This work |
| aBL416 | Δ*pyrE2* pTA962::halA-30aa-HaloTag pTA962::msfGFP-MTS-mYPet | This work |
| aBL420 | Δ*pyrE2* pTA962::halA-30aa-HaloTag pTA962::halA(Nt) | This work |
| aBL422 | Δ*pyrE2* pTA962::halA-30aa-HaloTag pTA962::halA(Nt)(ΔMTS) | This work |
| aBL439 | Δ*pyrE2* pTA962::halA-30aa-HaloTag | This work |
| aBL469 | *Δura3 ΔhalA* | This work |
| aBL564 | *ΔpyrE2 pyrE2::P250-msfGFP-mevR pTA962* | This work |
| aBL565 | *ΔpyrE2 pyrE2::P250-msfGFP-mevR pTA962::halA-40aa-msfGFP* | This work |
| aBL566 | *ΔpyrE2 pyrE2::P250-msfGFP-mevR pTA962::halA(Nt)(ΔMTS)-40aa-msfGFP* | This work |
| aBL567 | *ΔpyrE2 pyrE2::P250-msfGFP-mevR pTA962::halB-40aa-msfGFP* | This work |
| aKA11 | Δ*pyrE2* pTA962::HaloTag | This work |
| aZC36 | Δ*pyrE2* Δ*halB* | This work |
| aZC37 | Δ*trpA* Δ*pyrE2* Δ*halA* Δ*halB* | This work |
| aZC41 | Δ*halB* pTA962::halB-40aa-msfGFP | This work |
| aZC65 | Δ*trpA* Δ*pyrE2* Δ*halA* pTA962::ftsZ1-GFP | This work |

| | | |
|---|---|---|
| aZC67 | Δ*pyrE2* Δ*halB* pTA962::ftsZ1-GFP | This work |
| aZC69 | Δ*trpA* Δ*pyrE2* Δ*halA* Δ*halB* pTA962::ftsZ1-GFP] | This work |
| aZC71 | Δ*trpA* Δ*pyrE2* Δ*halA* ΔpyrE2 pTA962::ftsZ2-GFP | This work |
| aZC73 | Δ*pyrE2* ΔhalB pTA962::ftsZ2-GFP | This work |
| aZC75 | Δ*trpA* Δ*pyrE2* Δ*halA* Δ*halB* pTA962::ftsZ2-GFP | This work |
| aZC85 | Δ*trpA* Δ*pyrE2* Δ*halB* | This work |

**TABLE S5:** Oligos used in this work

| Alias | Sequence (5' → 3') |
|---|---|
| FW_up_halA | TATATTTCTAGATTCGAGCTTCCGGTCC |
| FW_down_halA | CGGTTTATTACGGCACGGCCGACGCCGAGGGTC |
| RV_up_halA | GACCCTCGGCGTCGGCCGTGCCGTAATAAACCG |
| RV_down_halA | TATATTAAGCTTTGGACGTCCGTCAGGC |
| oBL31 | AATTCGATATCAAGCTTATCGATTTCATTCATTTGTAAAGTTCATCCATTCCAT |
| oBL87 | CTTGTTTCGAGAGGGTTCAG |
| oBL97 | AGGTGGCACTTTTCGG |
| oBL105 | TGAGCAAAAGGCCAGC |
| oBL136 | CCGAAAAGTGCCACCTCGGCGGTAGAAGTACG |
| oBL137 | GCTGGCCTTTTGCTCAGTGGCGGATTCGATGTAG |
| oBL182 | CGGACCTATTGCGCATATGGTGTCACTGGGTACGG |
| oBL198 | GTCCGCTACCCTCAAGGGCCATCGGCGAGC |
| oBL318 | GGAATTCGATATCAAGCTTATCGATTTTTAGCCGCTGATTTCTAAGGTAG |
| oBL354 | CTTGAGGGTAGCGGAC |
| oBL355 | CGAGGAAGCGGAAGAGCG |
| oBL357 | GTCCGCTACCCTCAAGGATGACGATCCAGCCGT |
| oBL359 | CTGAGCCCGGTCCCTGGCCAGATCCCTCGAGGATGACGATCCAGCCGT |
| oBL360 | CCAGGGACCGGGCTCAGGCCAAGGTTCCGGCCCGCTCGTGATTCCAC |
| oBL369 | CCGAAAAGTGCCACCTGACGAGAAACGCGAGCA |
| oBL397 | CGGACCTATTGCGCATATGCTCTCCGGAGTCG |
| oBL426 | CGGACCTATTGCGCATATGTCGCCGCACGACC |
| oBL433 | GTCCGCTACCCTCAAGCAGTTTTGTTTTTTCTTTAATCTGTGAATG |
| oBL436 | CGCTGGTAATGAGGATACTGCGTGTCACTGGGTACGGAC |
| oBL440 | GCTCTAGAACTAGTGGATCCTTTACTCGGCTTCGCGAG |
| oBL441 | GCTGGTAATGAGGATACTGCATGTTCACAACGAAACAACTGG |
| oBL442 | GCTCTAGAACTAGTGGATCCTTCAGGCCATCGGCGAG |
| oBL472 | AGGAATTCGATATCAAGCTTATCGATTTTCAGGCCATCGGCGAG |

| | |
|---|---|
| oBL477 | GCACCGTACGTCTCGAGCAGAAGCTCGAACGCCT |
| oBL478 | GACGACCGTCAGACGAC |
| oBL479 | GTCGTCTGACGGTCGTCCGGACCATCAGGTTTCTTGA |
| oBL480 | ACGACGCATCCTGCAGCAAGCGCGATAGAGAGCAT |
| oHV36 | CGGACCTATTGCGCATATGGCAGAAATCGGTACTG |
| oHV67 | TACACGCTTGTGACTAATTCACT |
| oHV82 | CCCACTGCCTTGACC |
| oHV84 | GATACCGAAGCCGGGCGTGTTGGAGGACG |
| oHV186 | CGGACCTATTGCGCATATGGGAATTCTCTCTCGCA |
| oHV188 | GGTCAAGGCAGTGGGTCCGCCGGCGGGGCCGTCG |
| oHV189 | CAAGCTTATCGATTTCATTCACTCGGCTTCGCGAG |
| oJM160 | AGTGAATTAGTCACAAGCGTGTACATGGGAGGGGATGG |
| oJM212 | CTGAACCCTCTCGAAACAAGACCCGCCGACTCGGCGT |
| oJM213 | TCTTCCGCTTCCTCGTCATTTGTAAAGTTCATCCATTCCAT |
| oJM233 | AATCTATGGTCCTTGTTGGTG |
| oJM227 | TAACTGGCCGTCGTTTTAC |
| oJM228 | GGGCCCCTGGAACAG |
| oTR02 | GTCCGCTACCCTCAAGCTCGGCTTCGCGAG |
| oZC40 | GCCTCGGTGATTCCGGGAACCGGACGAACAATTATTATGG |
| oZC41 | GTTCGTCCACGAGGTC |
| oZC42 | GACCTCGTGGACGAACCGTGGATAAAACCCCTCG |
| oZC43 | GCTGGCCTTTTGCTCATTAGCCGTCGGCGTC |
| oZC44 | CCCGGAATCACCGAGG |

**Supplemental Movie Legends**

**Movie S1:** Halofilin mutants show different cell envelope defects. 3D-SoRa super-resolution projections of wild-type, Δ*halA*, Δ*halB*, and Δ*halAB* cells, as well as cells overexpressing *halA* and *halB*. Cells were stained with Nile Blue. Scale bar: 5μm.

**Movie S2:** HalA and HalB assemble, respectively, long filaments and foci, across the cytoplasmic membrane. 3D-SoRa super-resolution projections of rod (left) and disk (right) cells expressing HalA-msfGFP (orange) or HalB-msfGFP (blue). Cell membranes were stained with Nile Blue (grey). Scale bar: 2μm.

**Movie S3:** HalB-msfGFP foci are static over long periods. Cells were imaged at 2 minutes intervals using phase contrast and epifluorescence microscopy and displayed at 30 frames per second for 4.5 hours (approximately 2 generation periods). Scale bar: 5μm.

**Movie S4:** HalA-msfGFP filaments show a diversity of dynamics across the cell. Cells were imaged at 5 minutes intervals using phase contrast and epifluorescence microscopy and displayed at 7 frames per second for 6 hours (approximately 2.5 generation periods). Scale bar: 5μm.

**Movie S5:** Halofilin mutants have their cell shape impaired mildly during cytokinesis and cell elongation. Timelapses of cell division and elongation events of rod and disk cells were imaged at 2 minutes intervals using phase contrast microscopy and displayed at 12 frames per second for 3.6 hours (approximately 1.5 generation periods). Scale bar: 5μm.

**Movie S6:** Halofilin mutants have their cell shape compromised during cell shape transitions. Timelapses of cells shapeshifting from disks to rods were imaged at 2 minutes intervals using phase contrast microscopy and displayed at 15 frames per second for 5.3 hours (approximately 2.5 generation periods). Scale bar: 5μm.
